# Hypotrochoidal scaffolds for cartilage regeneration

**DOI:** 10.1101/2022.11.29.518172

**Authors:** Kenny A. van Kampen, Elena Olaret, Izabela-Cristina Stancu, Daniela F. Duarte Campos, Horst Fischer, Carlos Mota, Lorenzo Moroni

## Abstract

The main function of articular cartilage is to provide a low friction surface and protect the underlying subchondral bone. The extracellular matrix composition of articular cartilage mainly consists of glycosaminoglycans and collagen type II. Specifically the collagen type II organization has a characteristic organization in three distinct zones; (1) the superficial zone which has collagen fibers oriented parallel to the surface, (2) the intermediate zone where there is no predominant orientation, and (3) the deep zone which shows a high orientation with fibers perpendicular to the underlying bone. Collagen type II fibers in these 3 zones take an arch-like organization that can be mimicked with segments of a hypotrochoidal curve. In this study, a script was developed that allowed the fabrication of scaffolds with a hypotrochoidal design. This design was investigated and compared to a regular 0-90 woodpile design. The results showed that the hypotrochoidal design was successfully fabricated. Micro-CT analyses divided the areas of the scaffold in their distinct zones. In addition, the mechanical analyses revealed that the hypotrochoidal design had a lower component Young’s modulus while the toughness and strain at yield were higher compared to the woodpile design. Fatigue tests showed that the hypotrochoidal design lost more energy per cycle due to the damping effect of the unique microarchitecture. Finite element analyses revealed that the hypotrochoidal design had an improved stress distribution compared to the 0-90 woodpile design due to the lower component stiffness. In addition, data from cell culture under dynamic stimulation demonstrated that the collagen type II deposition was improved in the hypotrochoidal design. Finally, Alcian blue staining revealed that the areas where the stress was higher during the stimulation produced more glycosaminoglycans. Our results highlight a new and simple scaffold design based on hypotrochoidal curves that could be used for cartilage tissue engineering.

## 1. Introduction

Articular cartilage is an avascular tissue that is located in the joints [1]. The main function of cartilage is to protect the underlying subchondral bone from compressive loads and to provide a low-friction surface [2]. Injury or damage to cartilage can hamper this function and could lead to osteoarthritis [3]. Unfortunately, articular cartilage has a poor regenerative capacity due to the avascular and scarce cell density nature within the tissue [4]. Chondrocytes are the main cell population in articular cartilage and are responsible for synthesizing the extracellular matrix (ECM) [5]. The ECM of articular cartilage mainly consists of collagen type II, aggrecan and other proteoglycans [6]. The organization of the ECM is important in cartilage, specifically the collagen type II fibers have a particular organization which helps to improve the mechanical properties [7]. Cartilage is characterized by three distinct zones; (1) the superficial zone which has collagen fibers oriented parallel to the surface, (2) the intermediate zone where there is no predominant orientation and (3) the deep zone which shows a high orientation with fibers perpendicular to the underlying bone [8, 9]. Each of these zones have a different composition in ECM and cell density [10], which contributes to the unique properties of articular cartilage.

The organization of the collagen type II fibers could be mathematically described with arches or arch-like structures. A hypotrochoid is one of type of curve that could be used to describe an arch-like structure. This geometric curve that is generated by tracing a point that is linked to a smaller circle, which is rolling inside a larger circle [11, 12]. The hypotrochoidal curve is used to describe in a simplified way the satellite orbits around a planet in a solar system [13] and can be drawn using a tool called Spirograph [14]. The hypotrochoid is characterized by a closed outer surface with the outer shape being round in most cases. Depending on the parameters used, the hypotrochoidal curve can have a dense outer part with lines running parallel to the circle. These parallel lines, in combination with a more porous inner core with lines that run perpendicular to the surface of the circle, make it an interesting geometry that can be used to mimic cartilage ECM architecture. The generated arch-like structure resembles the collagen type II organization, especially if a segment on the top of the curve is taken. In addition, the complex arch-like structure should be able to distribute the forces through the whole scaffold more evenly [15, 16].

Additive manufacturing (AM) techniques such as fused deposition modeling (FDM) and bioprinting have been widely used to fabricate scaffolds for cartilage regeneration [17, 18]. The main advantage of AM techniques is the possibility to tailor the design and porosity of the scaffold [19]. Many scaffold fabrication approaches focus on matching the mechanical properties of native cartilage with a common strategy of changing geometry or by adding materials to the scaffold [20, 21]. While techniques such as FDM do pose the ability to produce complex shapes, in most cases a woodpile design is used where each layer is a composition of parallel running lines stacked in different angles with each layer [22]. Some efforts have been made to introduce complexity in the design, such as introducing gradients by increasing the spacing between the fibers in each layer [23, 24] or using different periodic infilling patterns [25]. However, none of these studies take a biomimicking approach from both a morphological and a mechanical point of view in the design of the scaffold. This could both be achieved by using a hypotrochoidal design.

In this study, we propose a new strategy to fabricate additive manufactured scaffolds with a hypotrochoidal pore network architecture. The mechanical properties of several hypotrochoidal designs were compared to a classical 0-90 woodpile structure. In addition, a finite element model was used to simulate the stress distribution through the scaffold. Finally, the effect of the hypotrochoidal design on cellular behavior was studied in both static and dynamic culture conditions.

## 2. Materials and methods

### 2.1 Scaffold fabrication

Poly(ε-caprolactone) (PCL) (Mn 45.000, Sigma-Aldrich,USA) was used to fabricated scaffolds via FDM using a Bioscaffolder (SysENG, Germany). Briefly, pellets of PCL were placed in a stainless steel syringe and heated up to 110 °C. A pressure of 4 bar was used to force the molten polymer to the extrusion screw, which was set at a constant rotational speed of 45 rpm. A 25G nozzle (260 μm internal diameter) was used for extrusion. The layer thickness and speed was kept constant at 180 μm and 400 mm/min, respectively.

A custom written python script (Python Software Foundation, Version 3.7.0) was developed to generate the deposition pattern for extrusion. Using a hypotrochoidal function both X and Y coordinates were calculated (Eq. 1 and Eq. 2 respectively). R = the radius of the large circle, r = the radius of the rolling circle and d = the distance between the center of the rolling circle and the point which is traced. Afterwards, cut off values were set to mark the boundaries of the scaffold. The hypotrochoidal scaffold followed the equation R = 10 and d = 4 with cutoff values between −5 and 5 in the X and bigger than 8 in the Y direction. As a variable parameter for the mechanical characterization, r was changed between 0.17 and 0.68. After each hypotrochoidal layer, two meandering layers that matched the outline of the scaffold were deposited with a fixed strand distance of 800 μm. The radius of this surface can be calculated with R = the radius of the large circle, r = the radius of the rolling circle and d = the distance between the center of the rolling circle and the point which is traced (Equation 3). As control, the hypotrochoidal layer was replaced with a meandering woodpile layer of similar tool pathway length but with a 90° rotation between layers (0-90).

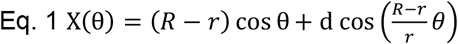

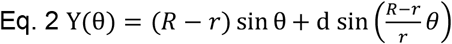

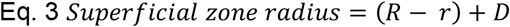

### 2.2 Scaffold characterization

Scaffold geometry and architecture was characterized by Stereomicroscopy (SMZ25, Nikon instruments) with a dark field illuminator (Nikon instruments). MicroCT was used to qualitatively and quantitatively investigate the 3D structure, volume and available surface area. The obtained micrographs were reconstructed using Nrecon software (Version 1.7.1.6.,Bruker MicroCT). The scanning parameters were set as follows: 2452×1640 camera resolution, 6.5 μm pixel size, source voltage and current of 50 kV and 130 μA respectively, 0.2 degree rotation step and 4 averaged frames. Pore shape was visualised using Ctvox (Version 3.3.0r1403, Bruker MicroCT) software and the quantitative data was analysed through Ctan software (1.18.4.0+, Bruker MicroCT). The scaffold was divided in three different sections based on the three zones found in articular cartilage [26], the deep, middle and superficial zone set at 25%, 55% and 20% of the scaffold, respectively. The fraction of the pore volume in each zone was derived from Eq. 4.

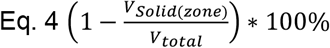

Where V_solid(zone)_ from each zone was obtained from the micro-CT analysis. V_total_ was calculated based on an equivalent solid block that was generated with computer-assisted design (CAD) (Rhino 6, Robert McNeel Associates, Version 6.19) with the dimension of the full scaffold.

### 2.3 Mechanical characterization

A custom designed adapter made of aluminum was used with a matching curvature of the scaffolds to maintain full contact area of the surface during mechanical testing (ElectroForce 3230, TA instruments, USA). Compression tests were performed using a 45 N load cell (ElectroForce) and the compression rate was set at 1% strain/s with a limit to 30%. A camera (DMC-G3, Panasonic) with macro lens (Panagor 90mm f2.8, Komine) was used for video acquisition during the compression test to observe the deformation. The Young’s Modulus was calculated from the linear part of the stress strain curves. The yield strain and strength were determined from the yield point that was set at the intersection between the stress-strain curve and the linear line drawn from the Young’s modulus with a 0.2% offset. The toughness of the scaffold was calculated as the area under the stress/strain curve up until the yield point. The maximum strain for the fatigue test was set at 2.5% based on the data obtained from the compression test. During that test 100 cycles were performed with a frequency of 1Hz. A single cycle of the hysteresis curve was plotted and the amount of energy lost per cycle was calculated as the difference in area under the curve between the loading and unloading part of the hysteresis. This was normalized against the total amount of energy to correct for differences between the scaffold designs.

### 2.4 Finite element modeling

A custom written script in Rhino grasshopper was used to convert the G-code of the hypotrochoidal design from the printer into a 3D CAD model. The CAD files of the tested designs were imported in finite element modeling software (COMSOL Multiphysics, COMSOL B.V. Version 6.0). The model was fixed at the bottom part and a concave fixture was made in the software to match the curvature of the scaffold. A strain of 10% was simulated in steps of 2% and the Von Mises stress along with the displacement were plotted.

### 2.5 Cell culture

ATDC5 (RIKEN cell bank), a teratocarcinoma derived chondrogenic cell line, was used to test the effect of the hypotrochoidal design on cells. Cells were trypsinized before reaching 80% confluence and 75.000 cells were seeded per scaffold. After 4 hours of attachment, the scaffolds were transferred into Dulbecco’s Modified Eagle Medium: nutrient mixture F-12 (DMEM-F12) (Sigma-Aldrich) expansion media supplemented with 5% fetal bovine serum (FBS) (Sigma-Aldrich) and 1% Penicillin Streptomycin (Thermo Fisher Scientific). After 1 day, the media was switched to differentiation media, which consisted of DMEM-F12 supplemented with 5% FBS, 1% Insuline, Transferin, sodium and Selenite (ITS) (Thermo Fisher Scientific), and 1% Penicillin Streptomycin. After 7 days, some of the scaffolds were transferred to a custom made bioreactor for dynamic stimulation. The stimulation was performed for 2 hours per day at 1Hz and 2.5% strain. The scaffolds were kept for a total of 28 days with media changes every other day.

### 2.6 Microscopy and cell analysis

Scaffolds were harvested after 28 days and used to analyze DNA, glycosaminoglycans (GAG) and collagen content or fixated for histological evaluation. A plate reader (CLARIOstar, BMG labtech) was used to quantify the biochemical assays. Scaffolds that were used for quantification were freeze/thawed 3 times before being submerged in 1 mg/ml proteinase K Tris/EDTA buffer solution. Samples were freeze/thawed again for three times after an overnight incubation at 56 °C in proteinase K solution. A part of the lysate solution was used to analyze DNA content through a DNA analysis kit (CyQUANT™ Cell Proliferation Assay Kit, Thermo Fischer Scientific) according to the manufacturer’s protocol. The samples were excited at 480 nm and emission was measured at 520 nm. The amount of DNA present in the sample was calculated from a known bacteriophage λ DNA standard. Another part of the solution was used to perform a GAG assay with 1,9-dimethylmethylene blue (DMMB) (Sigma-Aldrich) dye. The absorbance was measured both at 525 nm and 595 nm. The amount of GAGs in the sample was calculated by the difference in absorbance from both wavelengths and compared to a known chondroitin sulfate standard. The final part portion of the lysate was used to quantify the collagen content according to a hydroxyproline assay (Sigma-Aldrich). The absorbance was measured at 570 nm and the collagen content in the sample was determined from a hydroxyproline standard.

The scaffolds used for microscopy were fixed with 4% Paraformaldehyde (PFA) (VWR) for 1 hour and subsequently placed in PBS until further processing. Cells were stained with 1/250 dilution (0.23 μg/ml) DAPI (Sigma-Aldrich), 1/200 (0.5 μM) Phalloidin (Thermo Fisher Scientific), 1/400 dilution Collagen type II (Anti-Collagen type II, ab34712, Abcam) and 1/200 dilution Collagen type X (Anti-Collagen type X, ab49945, Abcam). The scaffolds were stained with an Alcian blue staining (Sigma-Aldrich) with a Nuclear red counterstaining (Sigma-Aldrich) to check the presence of GAGs. Fluorescent images were taken (Nikon Eclipse Ti-e) and the bright field images were acquired with the stereomicroscope (SMZ25, Nikon instruments).

### 2.7 Statistics

Statistical analyses were performed with GraphPad Prism 8.1.2. Significant differences were tested using an ANOVA test with a Dunnett multiple comparisons test and a student’s t-test to compare among two groups. The tests were considered significant when p<0.05.

## 3. Results

### 3.1 Scaffold fabrication and morphology

The nomenclature and fabrication parameters are shown in figure 1. Briefly, the full hypotrochoid was sectioned by boundaries to mimic the collagen type II alignment. The scaffolds were fabricated through FDM after digital sectioning. The final scaffolds were divided in three distinct zones based on literature, deep (25%), middle (55%) and superficial zone (20%) (supplementary information, Figure S1) [26]. For the hypotrochoidal designs, parameters R, d and the boundaries were kept the same while r varied between 0.17 and 0.68. As control a 0-90 design was compared to the hypotrochoidal designs.

**Figure 1.**
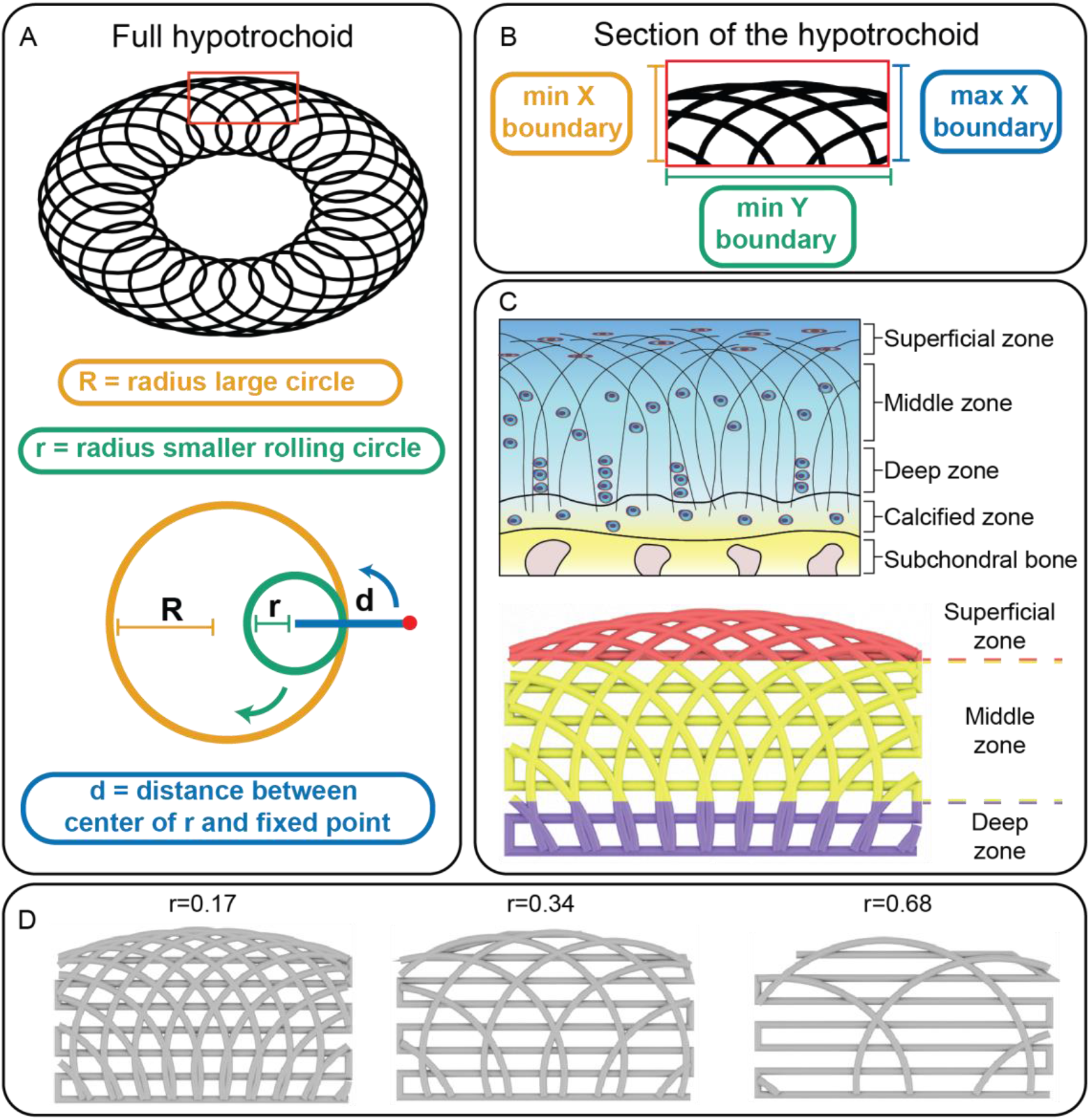
Nomenclature of the hypotrochoidal scaffold (A). Section of the hypotrochoidal scaffold (B). Different zones of the hypotrochoidal scaffold representing the different zones in articular cartilage, the fiber diameter in the scaffold is in the micron range while the bundles of collagen fibers is in the sub-micron range (C). Rendered images of the three tested designs with different r values (D).

As seen from both stereomicroscopy and micro-CT images the hypotrochoidal designs were successfully fabricated (Figure 2). The hypotrochoidal scaffolds had a curved top surface due to the way a hypotrochoid was generated. The meandering layers between the hypotrochoidal layers served as a support during the fabrication process. Signs of fiber collapse were noted in the designs that were more porous, due to the lack of support for the fabricated fiber that had to span in air multiple millimeters (Figure S2). The pores in all of the tested designs were interconnected. In the hypotrochoidal design, a gradient in pore size was observed with the smallest pores in the superficial zone and the largest pores in the deep zone. The volumetric analyses revealed that the scaffold volume of the hypotrochoidal design increased from 40.0 ± 0.4 mm^3^ to 65.0 ± 2.4 mm^3^ with a lower r value (r=0.68 and r=0.17 respectively) (table 1). A similar observation was made with the available surface area, increasing from 636.8 ± 14.5 mm^2^ to 919.3 ± 79.7 mm^2^ in the r=0.68 and r=0.17 samples respectively while the surface area to volume ratio remained similar between the hypotrochoidal conditions. The 0-90 structure was similar in volume to the r=0.68 scaffold (42.4 ± 1.2 mm^3^ and 40.0 ± 0.4 mm^3^ respectively) while having a higher surface area (722.0 ± 40.3 mm^2^ and 636.8 ± 14.5 mm^2^ respectively). The 0-90 structure, however, showed a higher surface area to volume ratio compared to the hypotrochoidal designs. The curvature of the different designs was shown in table 1 with minimal differences in radius between the scaffolds. The curvature for the 0-90 structure was set equal to the r=0.17 design.

**Figure 2.**
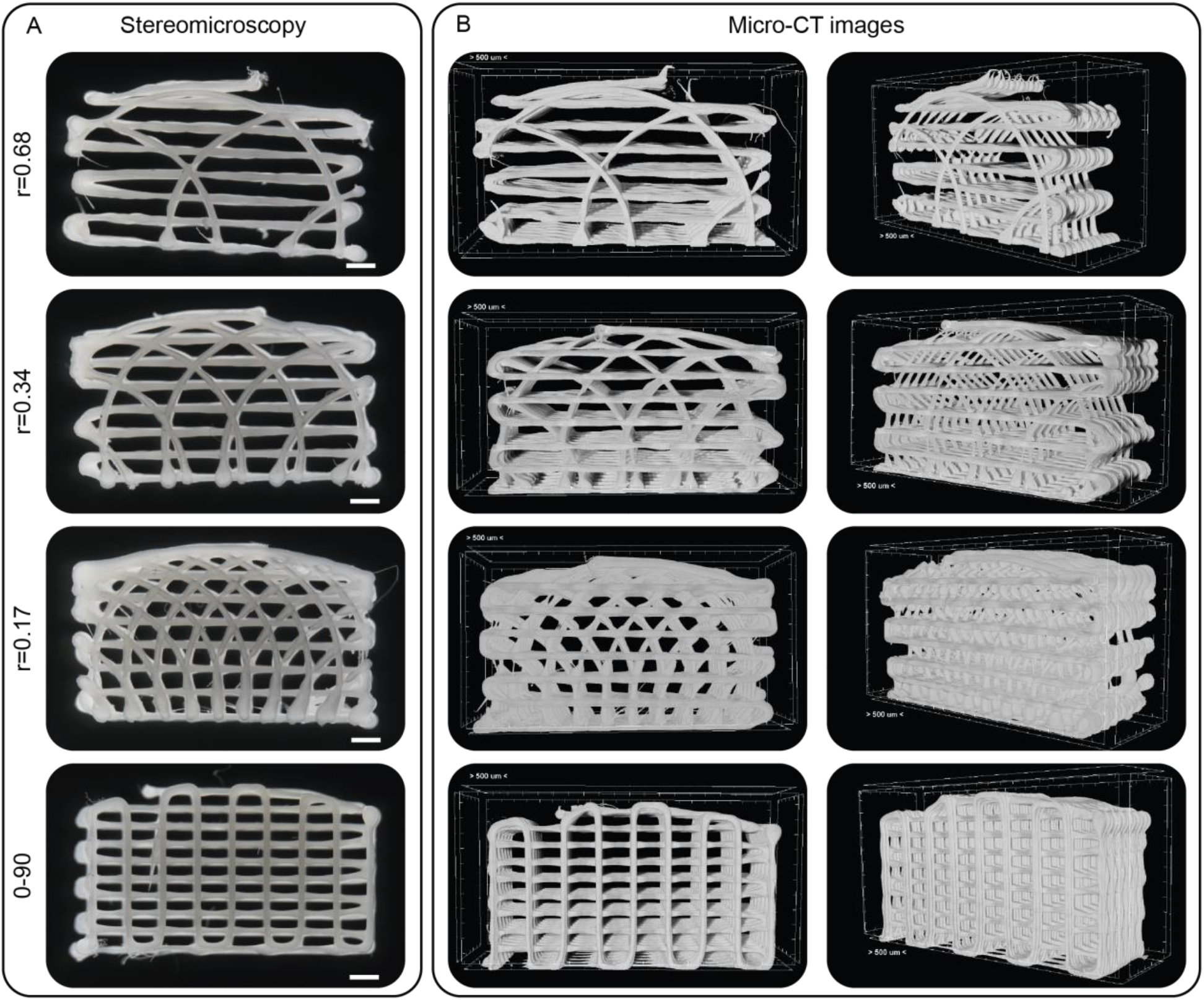
Overview of the tested designs. (A) Stereomicroscopy images of the frontal view of the tested designs. (B) The micro-CT images of the scaffolds, the frontal view in the left panels and the perspective view in the right panels. Scale bar represents 500 μm.

**Table 1.**
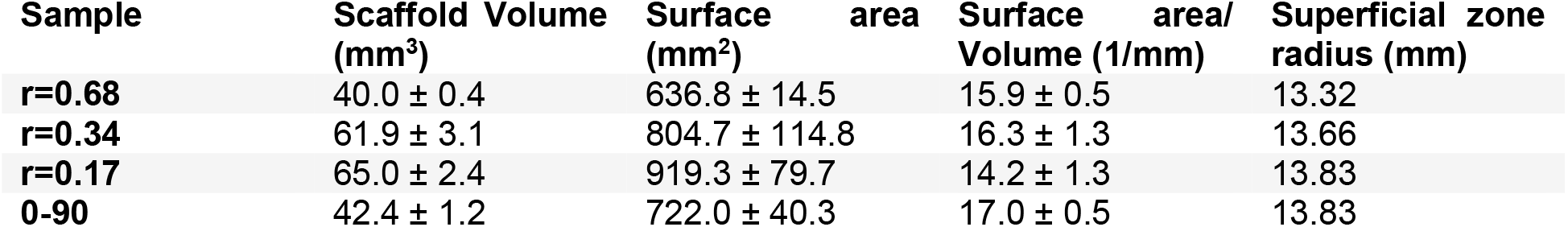
Volumetric micro-CT data of the different tested designs. Each condition contained n=3 samples. Values represent average ± standard deviation.

### 3.2 Mechanical testing and Finite element modeling

The mechanical properties of the hypotrochoidal designs were tested via compression and fatigue tests. CAD models of the scaffolds were also imported in finite element software and a compression test was simulated. The compression tests revealed that the Young’s modulus in the 0-90 structure was significantly higher compared to the r=0.17 hypotrochoidal designs (21.0 ± 1.8 MPa and 14.1 ± 1.8 MPa, respectively) (Figure 3A). In addition, the increase in r caused a decrease in Young’s modulus from 14.1 ± 1.8 MPa to 1.4 ± 0.1 MPa for r=0.17 and r=0.68 samples respectively. The differences in yield strain between hypotrochoidal designs was small ranging from 7.9 ± 0.0% and 11.0 ± 2.6% strain in the r=0.34 and r=0.68 respectively (figure 3B). The yield strain in the 0-90 design was significantly lower at 4.2 ± 0.4% compared to the hypotrochoidal designs. The yield strength in the hypotrochoidal designs decreased significantly with the increase in r from 36.9 ± 2.2 N to 5.9 ± 2.6 N (Figure 3C). The 0-90 design had a yield strength that was significantly different, between the r=0.17 and r=0.34 design (21.8 ± 3.6 N, 36.9 ± 2.2 N and 10.9 ± 0.6 N respectively). The toughness in the r=0.17 was significantly higher at 0.035 ± 0.001 N·mm^2^ compared to the other tested design at 0.008 ± 0.002 N·mm^2^, 0.011 ± 0.001 N·mm^2^ and 0.006 ± 0.004 N·mm^2^ for the 0-90, r=0.34 and r=0.68 design respectively (Figure 3D). The stress strain curves showed that the stress in 0-90 structure after the yield point decreased, while in all of the hypotrochoidal designs kept increasing (Figure 3E). Time lapsed snapshots showed that near the yield point 0-90 structure buckled in the middle of the scaffold and the square pore shape became rhomboidal (Figure 3F, supplemental video S1). However, the hypotrochoidal designs did not show this behavior. Instead, the top pores collapsed while the bottom pores remained intact, even far beyond the yield point (Figure S3). This was also confirmed during the compression test and micro-CT analyses where the 0-90 structure buckled in the top half of the scaffold, while the hypotrochoidal designs collapsed inward (Figure 4). Even at 25% strain the bottom part of the hypotrochoidal scaffolds showed interconnected pores (Figure S4).

**Figure 3.**
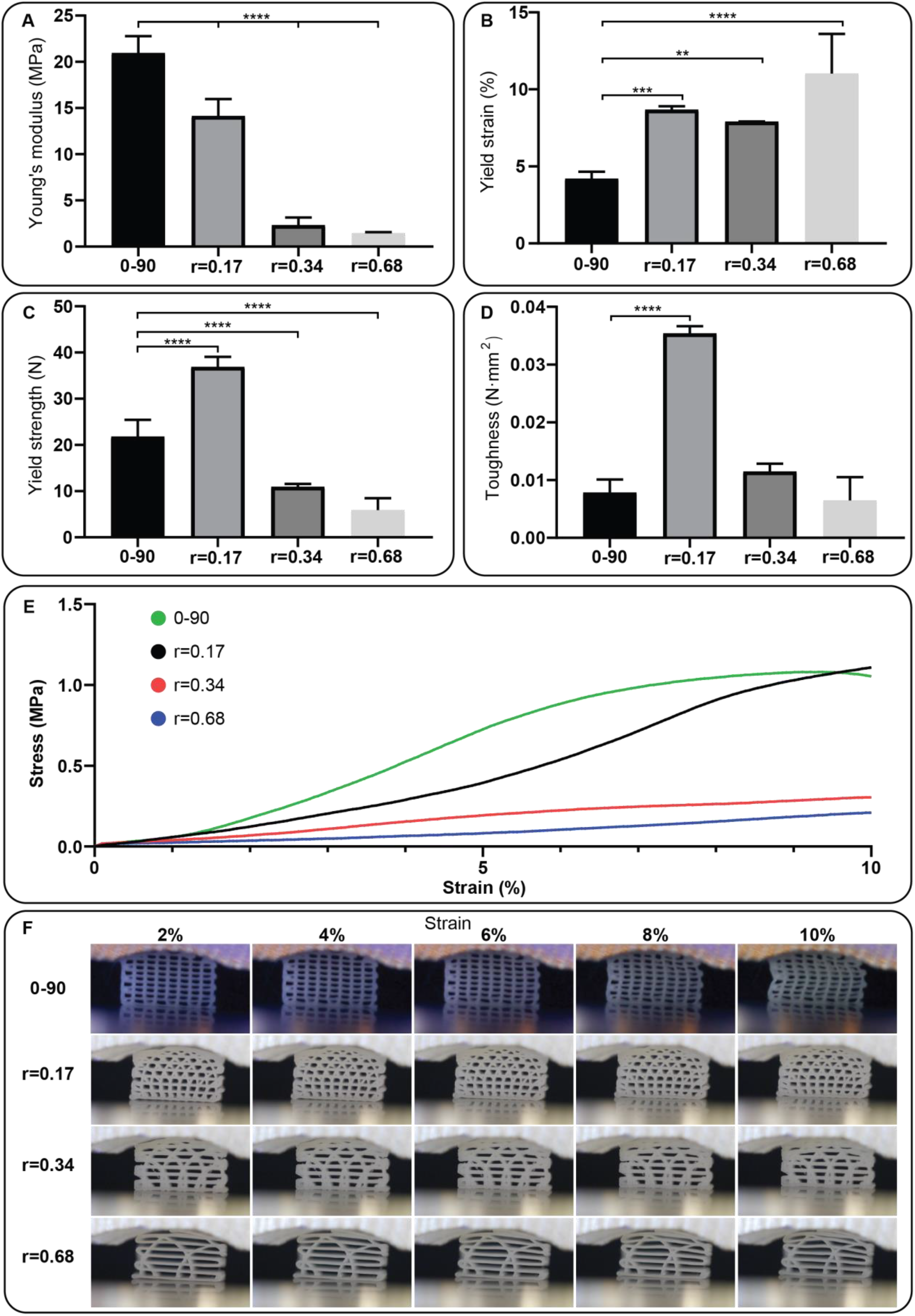
Compression test of the investigated designs: (A) Young’s modulus, (B) yield strain, (C) yield strength, (D) toughness, (E) representative stress strain curve, and (F) time-lapsed snapshots during the compression test. Each condition contained n=5 samples and values represent average ± standard deviation. Statistical significance: * p<0.05, **p<0.01, *** p<0.005, **** p<0.0001.

**Figure 4.**
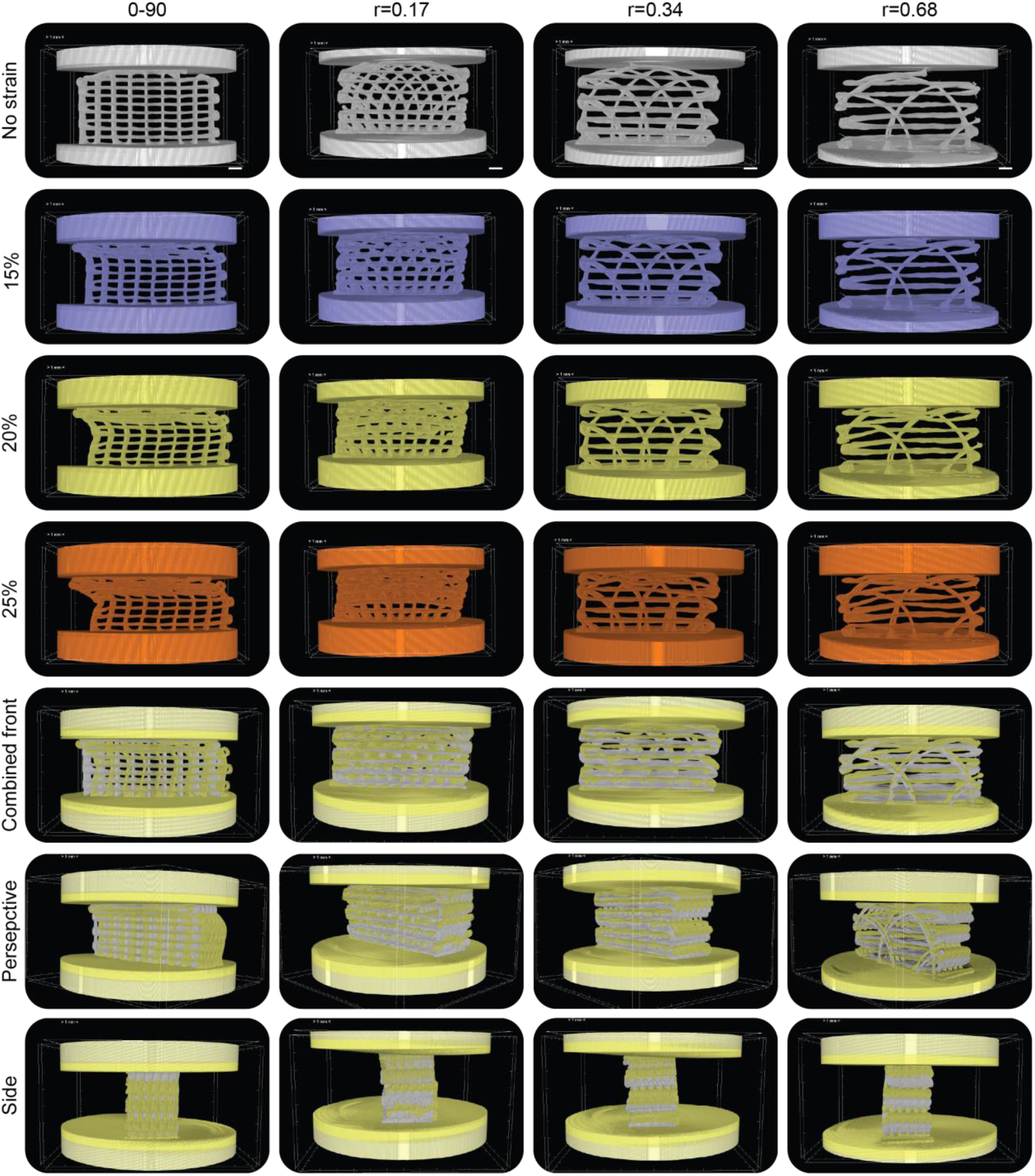
Micro-CT in combination with a compression test on the different tested designs. Images were taken at 15%, 20% and 25% strain. Panels from the bottom three rows are superimposed. Scale bar represents 500 μm.

A finite element model simulated up to 10% strain on the scaffolds with increments of 2% was performed. The model revealed that the vertical fibers in the 0-90 design endured the highest von Mises stress while the horizontal fibers were unaffected (Figure 5A). The displacement in the scaffold was evenly distributed from the bottom to the top of the scaffold. The stress distribution in the hypotrochoidal designs was improved compared to the 0-90 design, where the arches distribute the stress throughout the scaffolds and even in the meandering layer there was an increase in von Mises stresses. The highest stress was found in the superficial part of the scaffold (Figure 5B, C and D).

**Figure 5.**
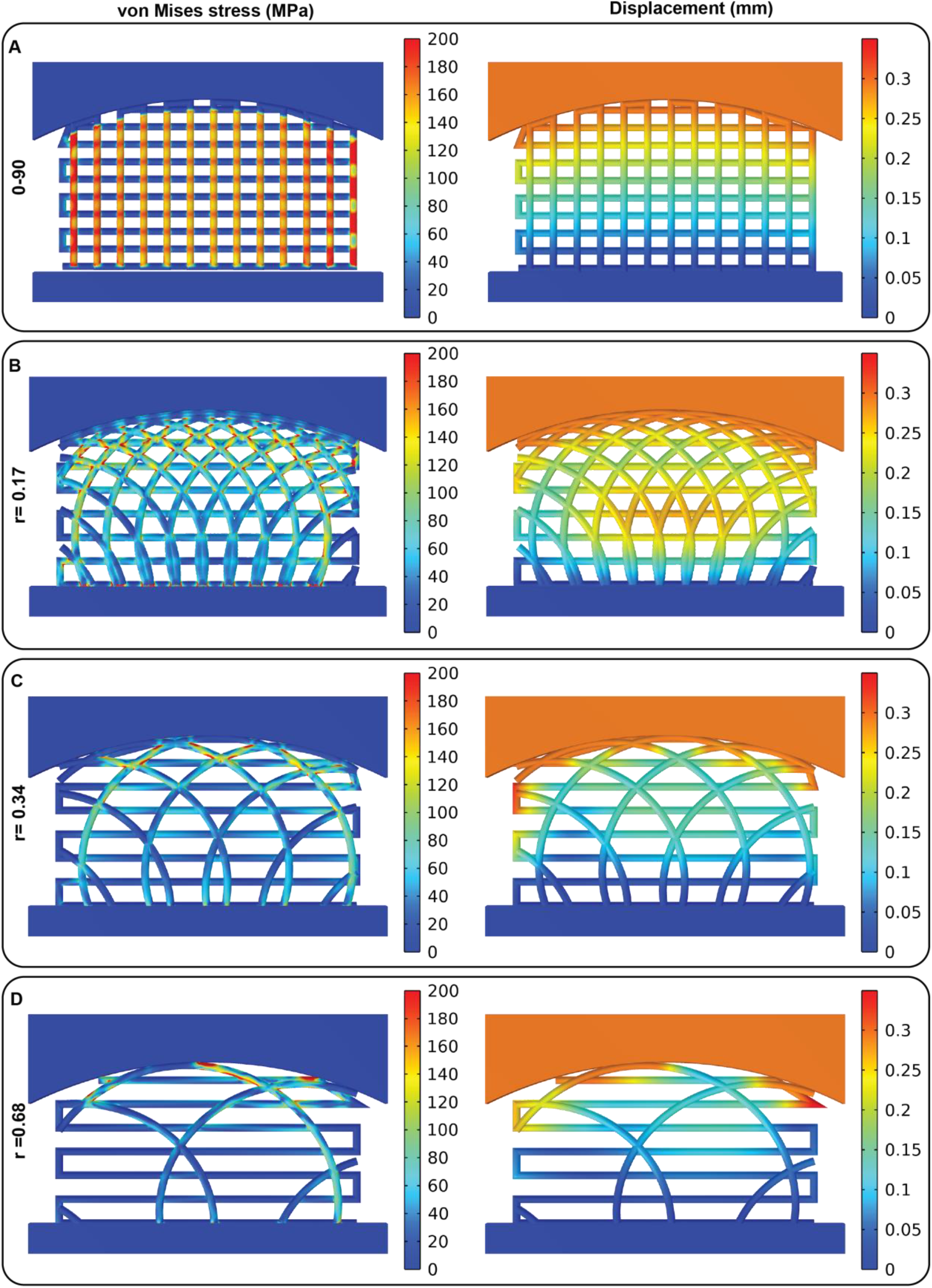
Finite element model of a compression test of the different designs, (A) the 0-90, (B) r=0.17, (C) r=0.34 and (D) r=0.68 design. The bottom fixture was fixed while a displacement of 0.6 mm was simulated on the curved top fixture (10% strain). The left hand panel show the von Mises stress and the right hand panel show the displacement.

Based on the results from the compression test, a fatigue test of 100 cycles at 1 Hz with a fixed 2.5% strain was performed. The data from the fatigue test showed that the hysteresis loop was the steepest in the 0-90 design compared to the hypotrochoidal designs (Figure 6A). The percentage energy lost per cycle, however, was significantly higher in the r=0.34 and the r=0.68 design compared to r=0.17 and 0-90 design (4.7 ± 0.5%, 6.0 ± 0.5% and 3.8 ± 0.5% respectively) (Figure 6B). The peak force in all of the samples decreased by a small amount in the initial cycles and stabilized after that initial decrease (Figure 6C). The average peak force in the 0-90 structure was the highest stabilizing around 12N while in the r=0.68 it was the lowest at around 2N.

**Figure 6.**
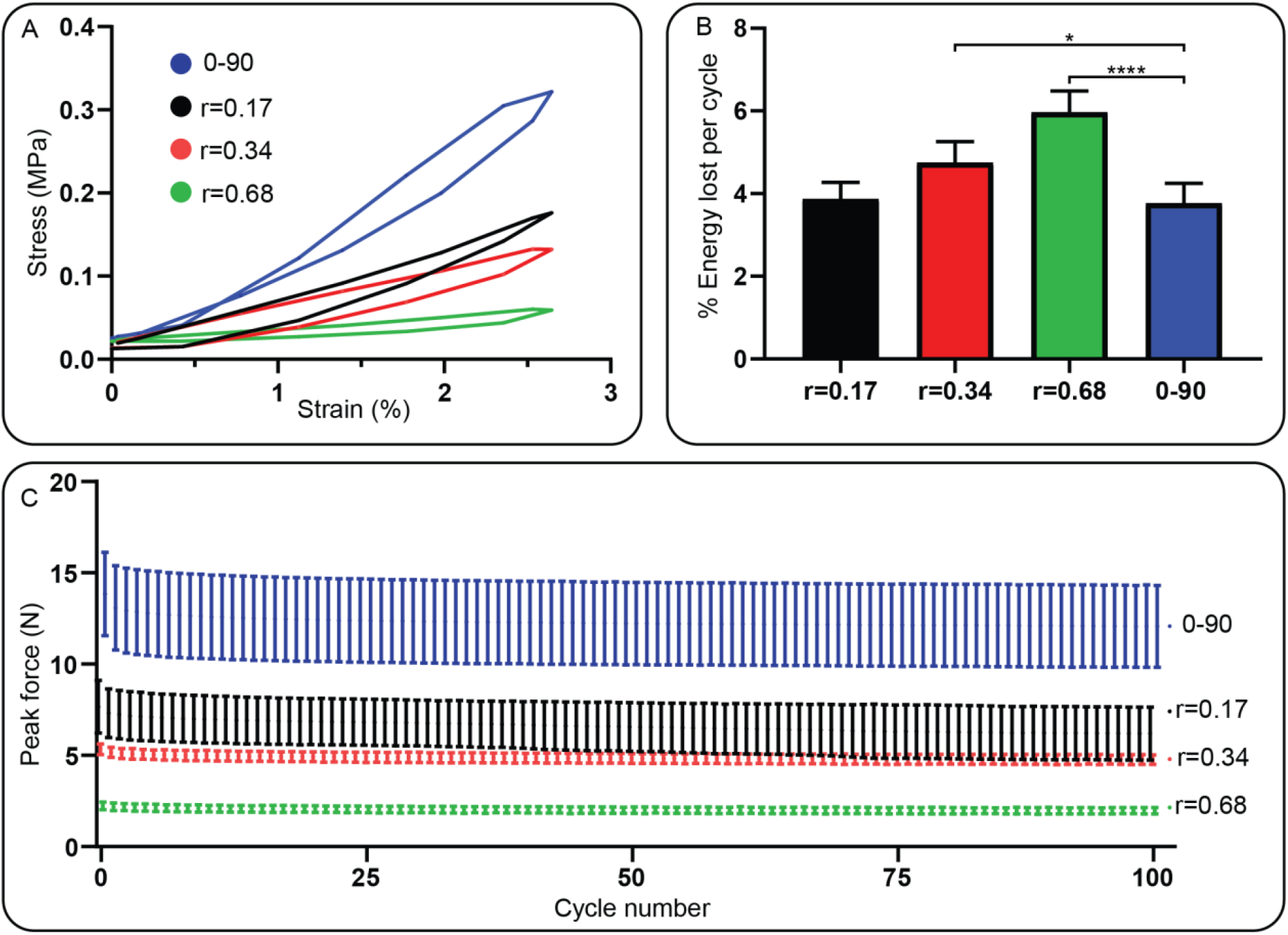
Fatigue test data with a frequency of 1 Hz until a strain of 2.5%. (A) Hysteresis loop of a single cycle during the fatigue test. (B) Average energy lost per cycle during the fatigue test. (C) Average peak force of each cycle. Each condition contained n=5 samples and values represent average ± standard deviation. Statistical significance: *p<0.05, **p<0.01, ***p<0.005, ****p<0.0001.

### 3.3 Dynamic cell culture

Cell culture experiments were performed on the r=0.17 design and the 0-90 woodpile structure to assess the cellular behavior on the hypotrochoidal design. The r=0.17 design was chosen since the pore size of the r=0.34 and the r=0.68 design were too large and would likely not be closed by sufficient regenerated tissue after the culture period. The fraction of the pore volume in both the 0-90 woodpile structure and the r=0.17 design was similar for each zone (Figure 7A). The scaffolds were seeded with ATDC5 cells and some of the scaffolds were transferred to a bioreactor after 7 days. A dynamic culture was performed by applying a mechanical stimulation for 2 hours every day at 1Hz and 2.5% strain until 28 days. The DNA in the samples was normalized against the pore volume. A division was made based on the fraction of the pore volume found in each zone with the assumption that the cells were equally distributed throughout the scaffold. The dynamic stimulation increased the DNA per pore volume in the r=0.17 samples from 2.9 ± 0.6 μg/mm^3^ to 3.7 ± 0.2 μg/mm^3^ respectively while there was a slight decrease observed in the 0-90 woodpile design from 3.9 ± 0.9 μg/mm^3^ to 3.7 ± 0.5 μg/mm^3^ (Figure 7B). Between the tested designs it was noted that the DNA per pore volume was approximately similar. The GAG/DNA per pore volume remained similar in all of the tested conditions (Figure 7C). Dynamic stimulation did increase the collagen/DNA per pore volume in the 0-90 woodpile design (4.7 ± 0.5 ng/μg/mm^3^ and 5.8 ± 0.4 ng/μg/mm^3^ respectively), while in the r=0.17 design it remained similar. Comparable values were observed if the values were corrected by the available surface area instead of the pore volume (Figure S5).

**Figure 7.**
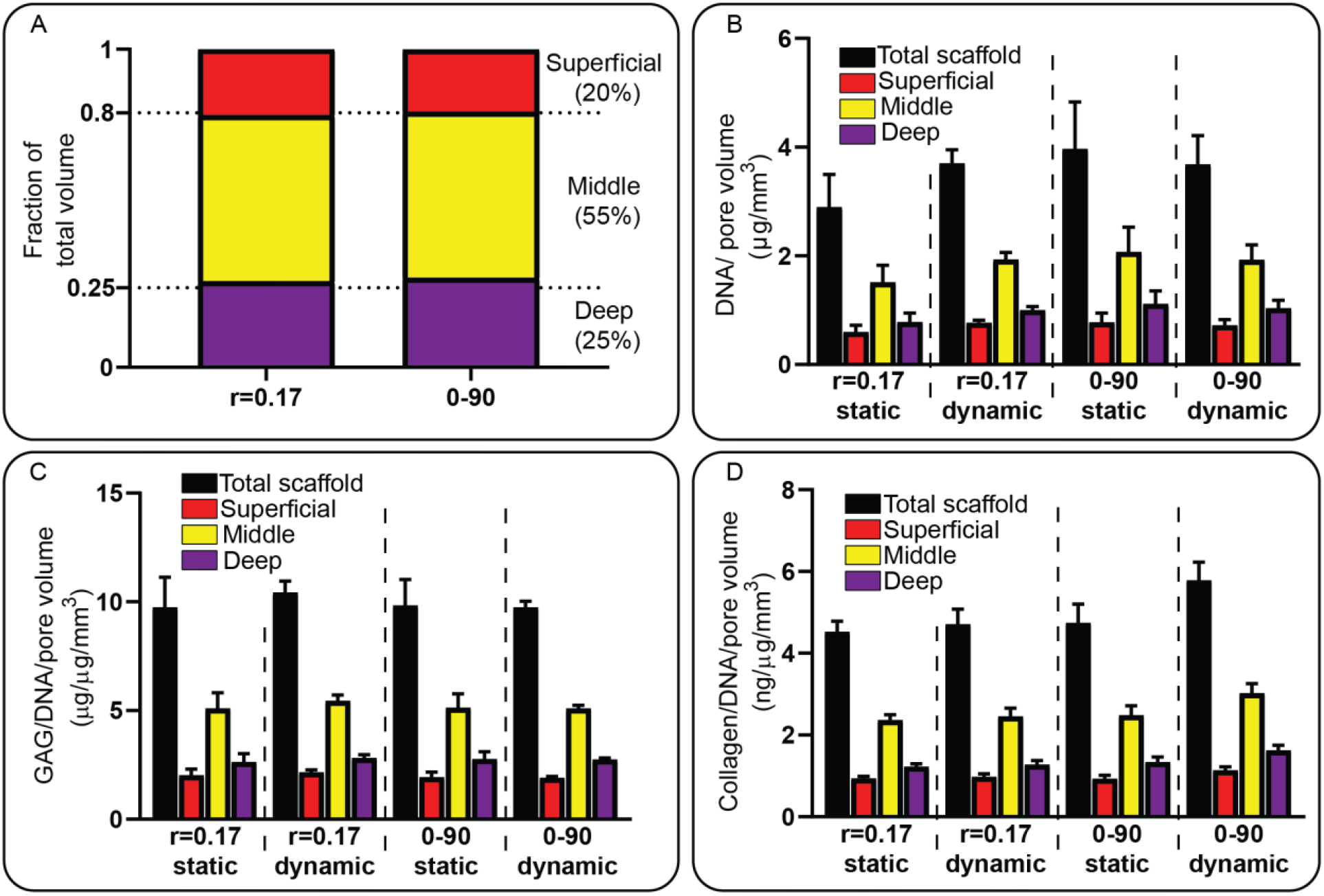
DNA, GAG and collagen analyses of the tested scaffolds after 28 days of culture. (A) Volumetric analyses of the pore volume as a fraction of the total volume. (B) DNA content normalized against the pore volume. (C) GAG per DNA content normalized against the pore volume. (D) Collagen per DNA content normalized against the pore volume. Each condition contained n=3 samples and values represent average ± standard deviation.

Alcian blue staining revealed that there was GAG deposition through the entire scaffold in all of the conditions (Figure 8). More pores in the 0-90 woodpile static condition seemed to be open compared to the dynamically stimulated samples. The intensity of the staining however was stronger in the static samples. The majority of the pores in r=0.17 hypotrochoidal design were open in both the static and dynamic samples, except for the smaller pores that were mainly located in the superficial zone of the scaffold. The r=0.17 dynamically stimulated samples showed darker areas with more GAG deposition that correlated with areas that should have received less stress during the culture located in the deep zone of the scaffold.

**Figure 8.**
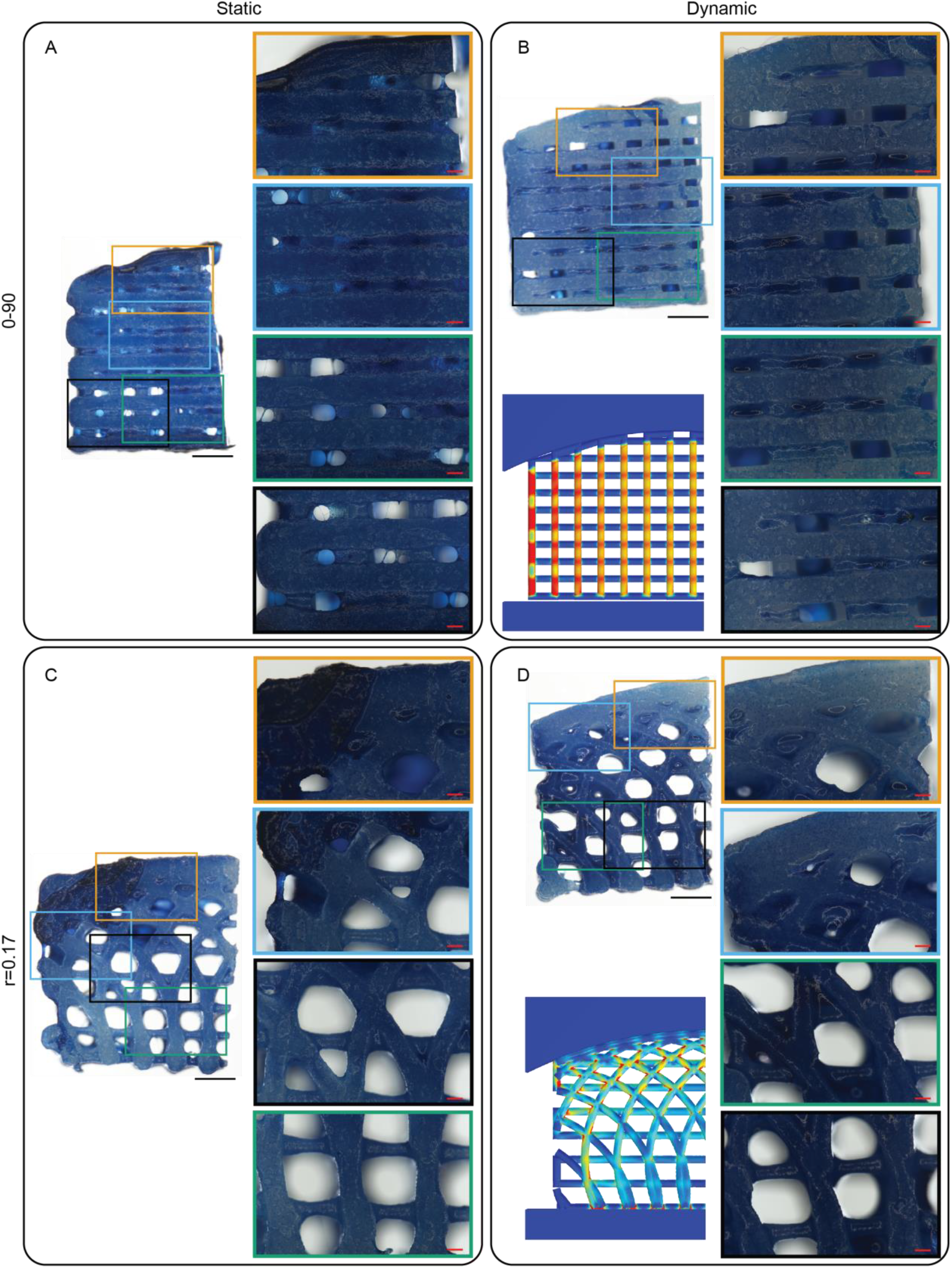
Alcian blue staining of the tested scaffold after 28 days of culture. The left panels show a stained scaffold and in the right panels a close up of selected areas. A finite element model with the von Mises stress as a reference in the dynamic samples. (A) 0-90 woodpile design after static culture. (B) 0-90 woodpile design after dynamic culture. (C) r=0.17 hypotrochoidal design after static culture. (D) r=0.17 hypotrochoidal design after dynamic culture. Scale bar in the left panels represents 1 mm; in the close up panels 200 μm.

Fluorescent images from the dynamic culture revealed collagen type X expression in the 0-90 woodpile design. Combining the data from the finite element model revealed that in the areas where there was more von Mises stress the amount of collagen type X expression was higher. There was almost no collagen type II expression in the 0-90 woodpile design. In the hypotrochoidal design, higher collagen type II expression was observed in the areas where the von Mises stress was increased. In the static conditions there was no collagen type II expression observed (Figure S6). The expression of collagen type X was comparable throughout the scaffold.

**Figure 9.**
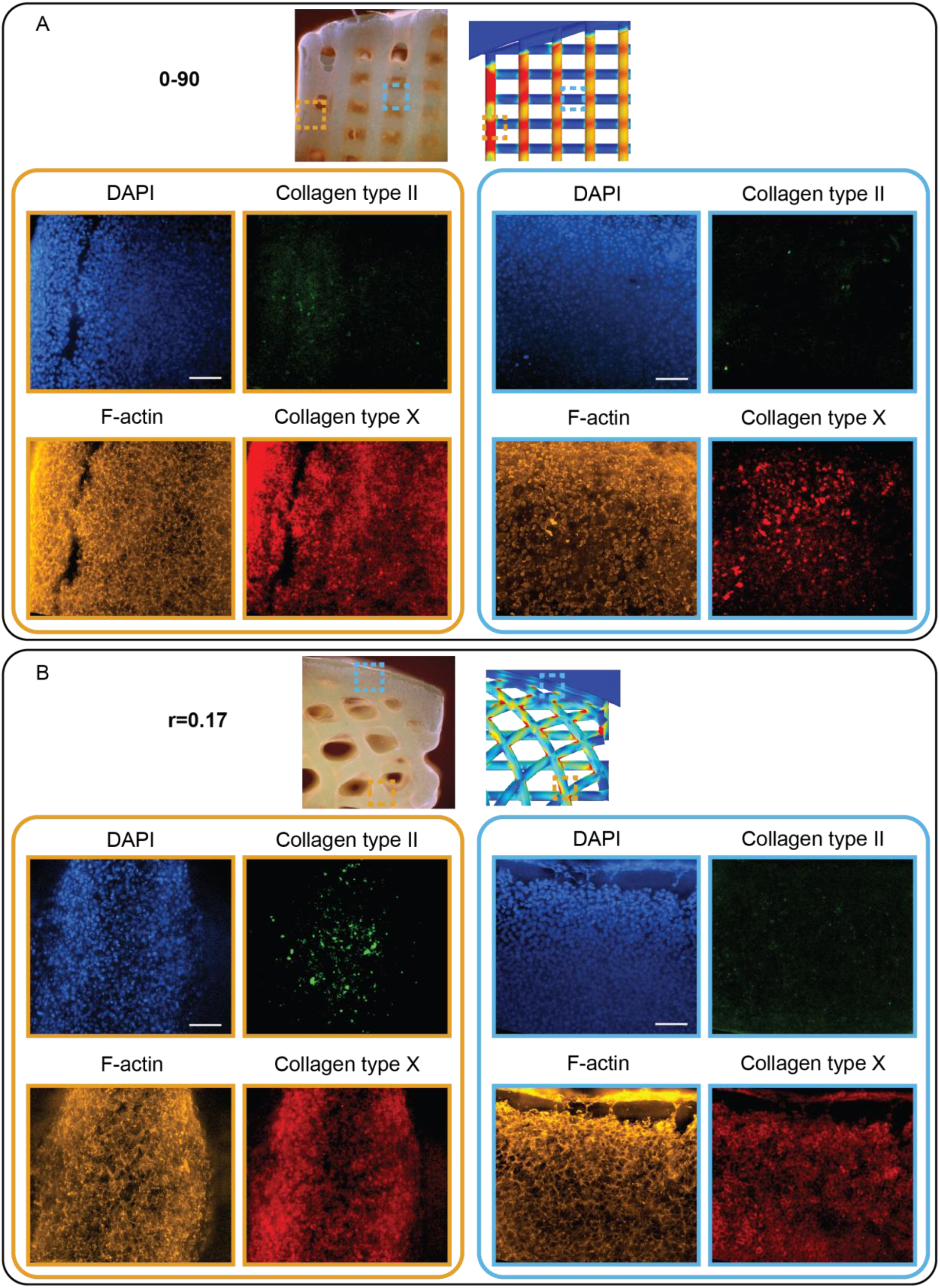
Cells after 28 days in dynamic culture. On top of the panel an overview image of the scaffold with the finite element model as reference. The left panel set with orange boundaries an area with a high amount of von Mises stress and the right panel set with blue boundaries an area with a low amount of von Mises stress. (A) 0-90 woodpile design. (B) r=0.17 hypotrochoidal design. Scale bars represent 500 μm in the overview panel and 200 μm in the close up.

## 4. Discussion

In this study, a scaffold was created from a hypotrochoidal deposition pattern. The morphology of the hypotrochoidal scaffold resembled that of the collagen fiber architecture in articular cartilage [8]. In addition, micro-CT analyses revealed that the hypotrochoidal scaffold had a fully interconnected pore network, which is necessary to have a proper access to nutrients [27]. The mechanical tests showed that the hypotrochoidal design had a lower Young’s modulus while having a significantly higher yield strain and higher toughness. In addition, the finite element model demonstrated that the stress distribution was improved with the hypotrochoidal design compared to the 0-90 woodpile structure. Further mechanical analysis from the fatigue test revealed that the hypotrochoidal design dissipated more energy per compression cycle. Finally, the dynamic culture demonstrated that the hypotrochoidal design had improved collagen type II deposition. Hence, the hypotrochoidal design could potentially be used for cartilage tissue regeneration.

The micro-CT analysis revealed that there was a pore gradient in the hypotrochoidal design, where the smallest pores resided in the top part of the scaffold and the largest pores in the bottom. This gradient can be beneficial since the structure of articular cartilage is heterogeneous, resulting in different regions of cartilage having a different range in mechanical properties [10]. Due to how a hypotrochoid is generated, the superficial zone of the scaffold always has a curved surface. The surface of healthy articular cartilage, however, is not perfectly round and is almost flat [28]. A solution to this could be to have R at least a factor 1000 bigger compared to r. This would approximate a flat surface while still creating a hypotrochoidal design. The chondrocytes in the superficial zone are responsible for the synthesis of superficial zone protein that acts as a lubricant [29]. As the hypotrochoidal curve is a closed curve, the hypotrochoidal design always lead to a non-porous surface. Therefore adding a single layer of a woodpile structure could be a strategy to introduce some porosity in the surface, which may facilitate the migration of cells to the surface of the scaffold to secrete superficial zone protein.

It was noted that the surface area to volume ratio in the hypotrochoidal design was lower compared to the 0-90 woodpile structure. An explanation for this could be that the fibers in the 0-90 design have the least amount of overlap in fibers due to a fiber angle of 90°, while the fiber angle in the hypotrochoidal design vary with every fiber. Having a lower surface area to volume ratio could interfere with transportation of nutrients and vasculature [30]. However, for cartilage this could be less important since it is avascular and relies on nutrients through diffusion [31].

The mechanical properties for a tissue such as cartilage are important [32]. The Young’s modulus of articular cartilage ranges from 10.6 – 18.6 MPa in the ankle joint to 5,5 – 11.8 MPa in the knee [33]. Specifically the r=0.17 design resulted within this range with an average of 14.1 MPa while the 0-90 woodpile design was slightly above the maximum range with 21 MPa. Even though the 0-90 woodpile structure is stiffer, it can absorb less energy before permanently deforming as was shown by the compression test. The contribution of the arches in the hypotrochoidal design could be the reason why the 0-90 woodpile design is less tough and stiffer. The arch architecture of the hypotrochoidal design can distribute the force through the scaffolds, as was shown in the finite element models. Similar findings from Chen et al. showed that a pillar design caused a higher stress more focused on the pillars compared to an octet design with more curved features [34]. The arch architecture in the hypotrochoidal design also showed that the collapse behavior was different. While the vertical fibers in the 0-90 woodpile structure buckled, the hypotrochoidal scaffold collapsed gradually, with the superficial zone collapsing before the deeper zone. Similar to articular cartilage, after impact the superficial zone is the first zone to receive permanent damage [35]. Besides compressive forces, articular cartilage is also subjected to shear forces [36]. The hypotrochoidal design could prove beneficial as it is known that arches can distribute the shear forces through the entire structure [37]. Another observation that was made is that the energy lost per cycle in the fatigue test was higher in the r=0.34 and r=0.68 hypotrochoidal design compared to the 0-90 woodpile structure, indicating that having less arches in the scaffold resulted in a higher energy loss. It is reported that articular cartilage behaves as an elastic material during hysteresis, with a relative energy loss of 28% [38]. The same study also found that a higher energy loss is also linked to cartilage damage. All the tested designs were below 10% energy loss which is well below the relative energy loss in articular cartilage. This value can change however, depending on the magnitude of the strain applied [39, 40].

GAG content is important for cartilage as it retains the water in the cartilage that can act as a lubricant [41, 42]. Our results showed that the GAG content per DNA remained similar in all of the conditions. It is important to note that the GAG content from the biochemical assay is performed on the entire scaffold, averaging the results from all the zones. The division of the zones was based on the micro-CT analyses with the assumption that the cells were distributed equally through the scaffolds. However, it could be that local areas that experienced more stress can contain higher amounts of GAG. This was demonstrated by histological analysis where certain areas were darker for Alcian Blue staining, thus possibly containing more GAGs. Specifically in the deeper zone of the hypotrochoidal scaffold, the intensity of the Alcian blue staining was higher. This correlates with a native hyaline cartilage tissue where GAG content is higher in the deeper zone compared to the middle and superficial zone [10, 43], Possibly due to GAG content being linked to improved mechanical properties and its deposition increased during mechanical loading [44]. Similar observations could be made for collagen content quantification, as also the collagen content is different for each zone with the highest amount of collagen found in the superficial surface [5]. The total collagen content was increased with the dynamic stimulation in the 0-90 woodpile structure. While the dynamic stimulation had no effect on the hypotrochoidal design. Examining immunostaining, however, revealed that the collagen that was deposited in the 0-90 woodpile design was collagen type X, which is a known hypertrophic marker [45]. ATDC5 cells are known to be hypertrophic and deposit collagen type X [46]. A study by Shukunami et al. shows that collagen type X deposition starts after 21 days in 2D [47]. In this study, the collagen type X deposition was primarily deposited in the areas where the von Mises stress was the highest according to the finite element model. The fibers within the samples that were unstimulated had a lower collagen type X deposition. Interestingly, collagen type II expression was increased in the hypotrochoidal design, specifically in the areas where the von Mises stress were high. The von Mises stress in the hypotrochoidal design were not as high as in the 0-90 structure indicating that collagen type II deposition can be enhanced by modulating the von Mises stress. Another method to increase collagen type II deposition for this study is to extend the dynamic stimulation until 4 weeks, as was shown with primary chondrocytes [48]. Culturing the scaffolds under a lower oxygen concentration in combination with the dynamic stimulation could be an additional step that might be considered in future studies, since a lower oxygen concentration is known to improve collagen type II expression [49, 50].

Generally, bundles of collagen fibers have a diameter ranging from 0.7 to 5 μm [51, 52]. The diameter of the fibers produced in this study are two orders of magnitude bigger. Therefore, mimicking the collagen fiber arrangement could prove challenging with a technique such as FDM. However, the hypotrochoidal design could serve as a blueprint for collagen fiber deposition as collagen is synthesized intra-cellular and assembled extracellular into fibrils by chondrocytes [53]. The cells that attach to the fibers of the scaffold could start secreting pro-collagen and assemble it into collagen fibrils following the hypotrochoidal pattern. Other techniques such as single cell acoustic patterning already allow cells to be arranged in a way that mimics the deep zone of cartilage [54]. The combination of cell patterning and having a template for the cells to follow could be worth exploring. The manufacturing of hypotrochoidal design scaffolds is not limited to FDM systems. Since the equations to generate X and Y coordinates are known, any system that uses an XY gantry could fabricate hypotrochoidal designs. Some alternative methods to produce these scaffolds might be with bioprinting or melt electrowriting systems (supplemental Figure 7). Specifically with melt electrowriting smaller fiber diameter can be obtained reaching the diameter of the collagen fiber bundles [55]. The hypotrochoidal design could potentially be used for other tissues where a gradient occurs such as long bones [56], due to the empty inner circle in the hypotrochoid that resembles the medullary cavity and the dense outer structure resembling the compact bone [57] (supplemental Figure 7).

## 5. Conclusion

Here, we showcased a hypotrochoidal design for cartilage tissue engineering. Such scaffolds mimic the morphological architecture of collagen fibers in articular cartilage, despite being still an order of magnitude larger than native collagen. It was shown that the hypotrochoidal design had an improved toughness compared to a 0-90 woodpile design. In addition, the stress distribution in the hypotrochoidal was improved due to the arched architecture that was introduced. The dynamic culture showed that the collagen deposition was improved and the hypotrochoidal design showed synthesis of collagen type II deposition in the specific areas of the scaffolds with a higher amount of stress.

## Abbreviations

ECM: extracellular matrix
AM: Additive manufacturing
FDM: Fused deposition modelling
PCL: Poly(ε-caprolactone)
CAD: computer assisted design
GAG: glycosaminoglycan

## CRediT authorship contribution statement

**Kenny A. van Kampen:** Conceptualization, Methodology, Formal analysis, Investigation, Writing – Original Draft, Visualization

**Elena Olaret**: Formal analysis, Investigation, Writing – Review & Editing, Visualization

**Daniela Campos:** Resources, Writing – Review & Editing, Supervision

**Horst Fischer:** Resources, Writing – Review & Editing, Supervision

**Izabela-Cristina Stancu**: Resources, Writing – Review & Editing, Supervision

**Carlos Mota:** Conceptualization, Resources, Writing – Review & Editing, Supervision

**Lorenzo Moroni:** Conceptualization, Resources, Writing – Review & Editing, Supervision, Funding acquisition

## Acknowledgements

This work was supported by H2020 FAST (NMP-7, GA n. 685825), the ERC Cell Hybridge (GA n. 637308). The microCT analyses was supported by the European Regional Development Fund (6695072), through Competitiveness Operational Program 2014-2020, Priority axis 1, ID P_36_611, MySMIS code 107066, INOVABIOMED, Romania. We also gratefully acknowledge Dr. J. Fernández-Pérez for the supervision during the hydroxyproline assay, A. Chandrakar for the melt electro written scaffold, Dr. M. Decarli for the bioprinting of the hypotrochoidal scaffold.

## Supplementary material

**Figure S1.**
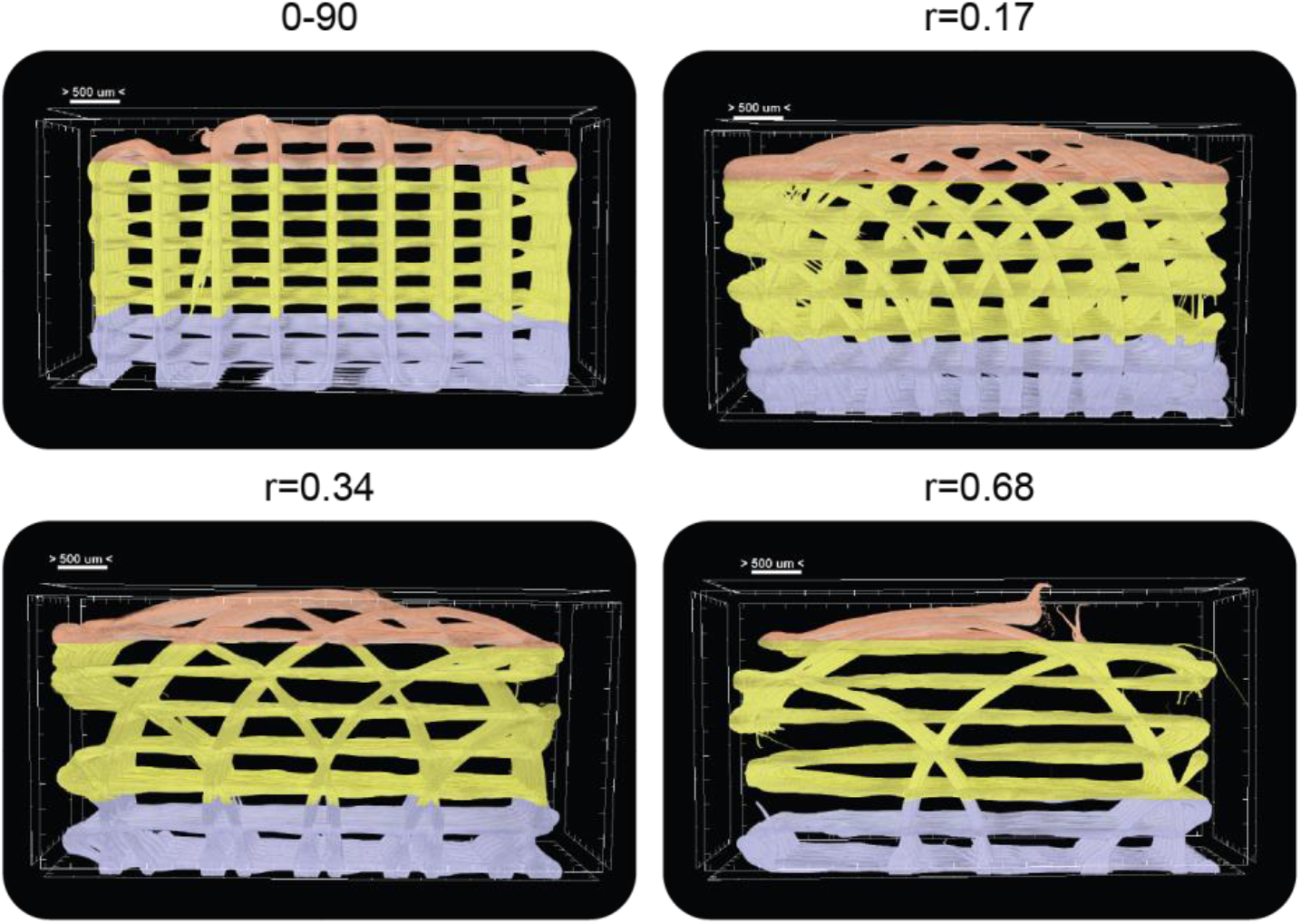
Division of the different scaffold designs into the three zones. The deep zone in blue, middle zone in yellow and superficial zone in red. Scale bar represents 200 μm.

**Figure S2.**
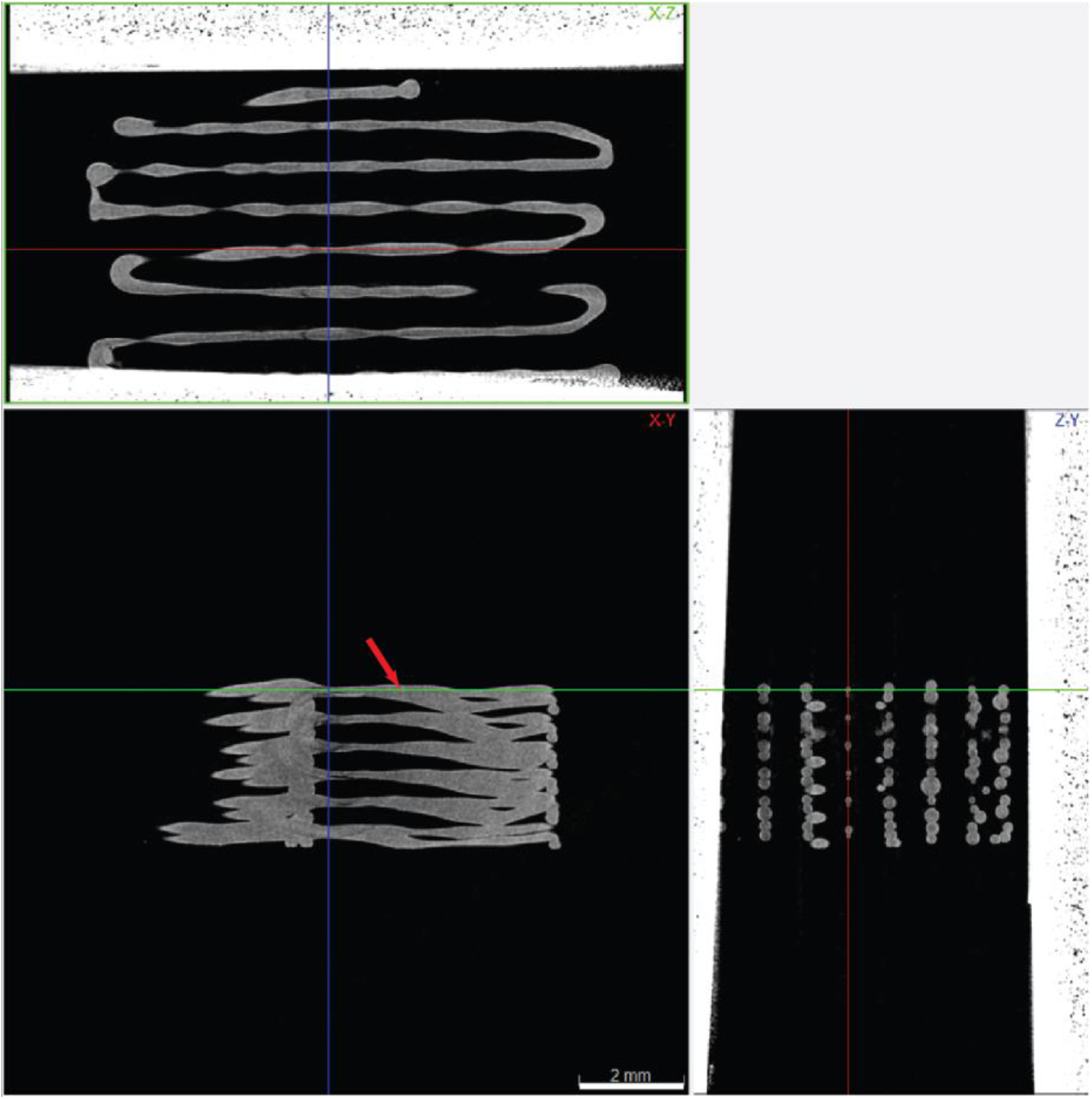
Fiber collapse in the r=0.68 hypotrochoidal design. Top panel displays the frontal view, the bottom left panel top view and the bottom right panel the cross sectional view. The red arrow indicates the collapse of the fiber. The scale bar represents 2 mm.

**Figure S3.**
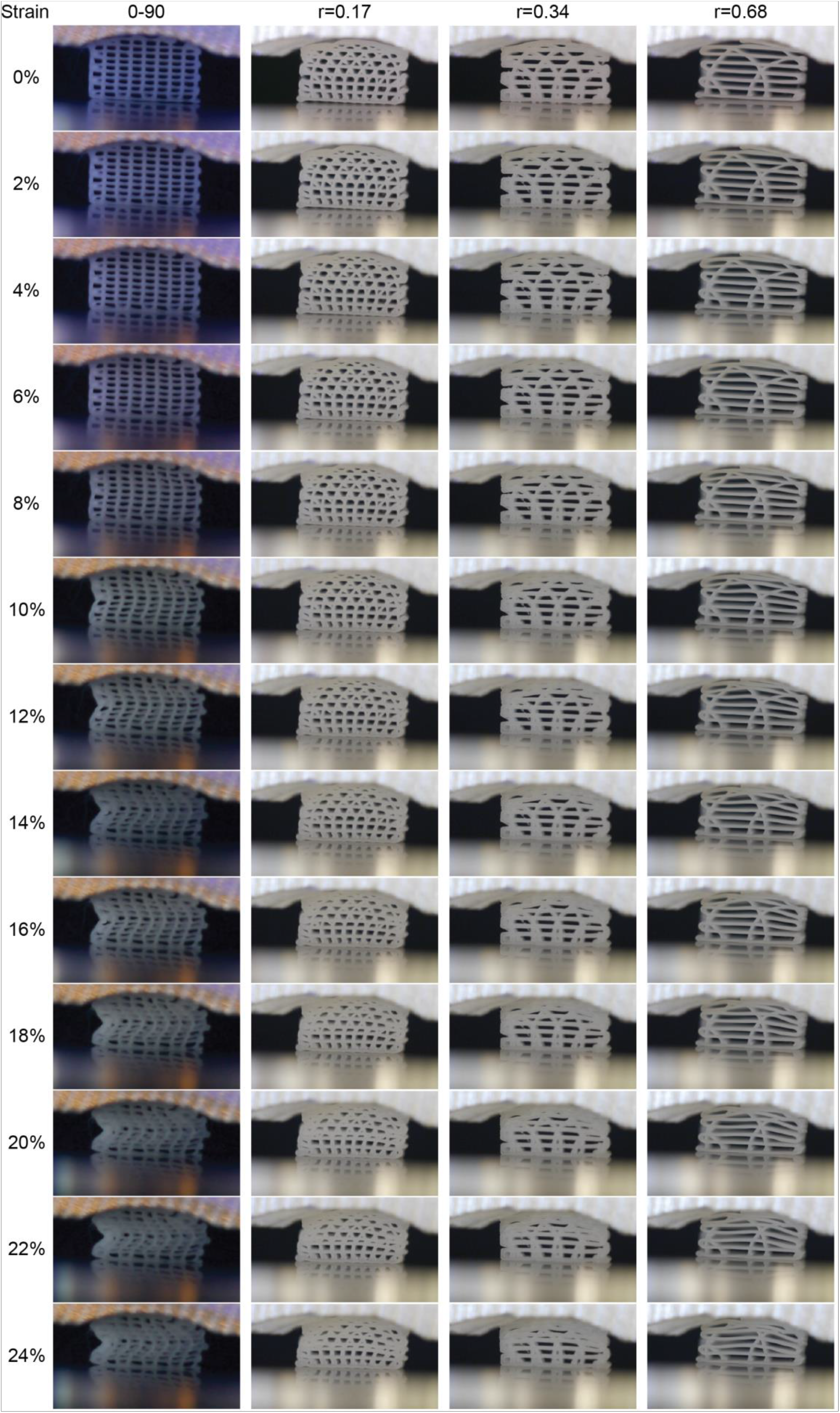
Time-lapsed snapshots during the compression test of the different tested designs.

**Figure S4.**
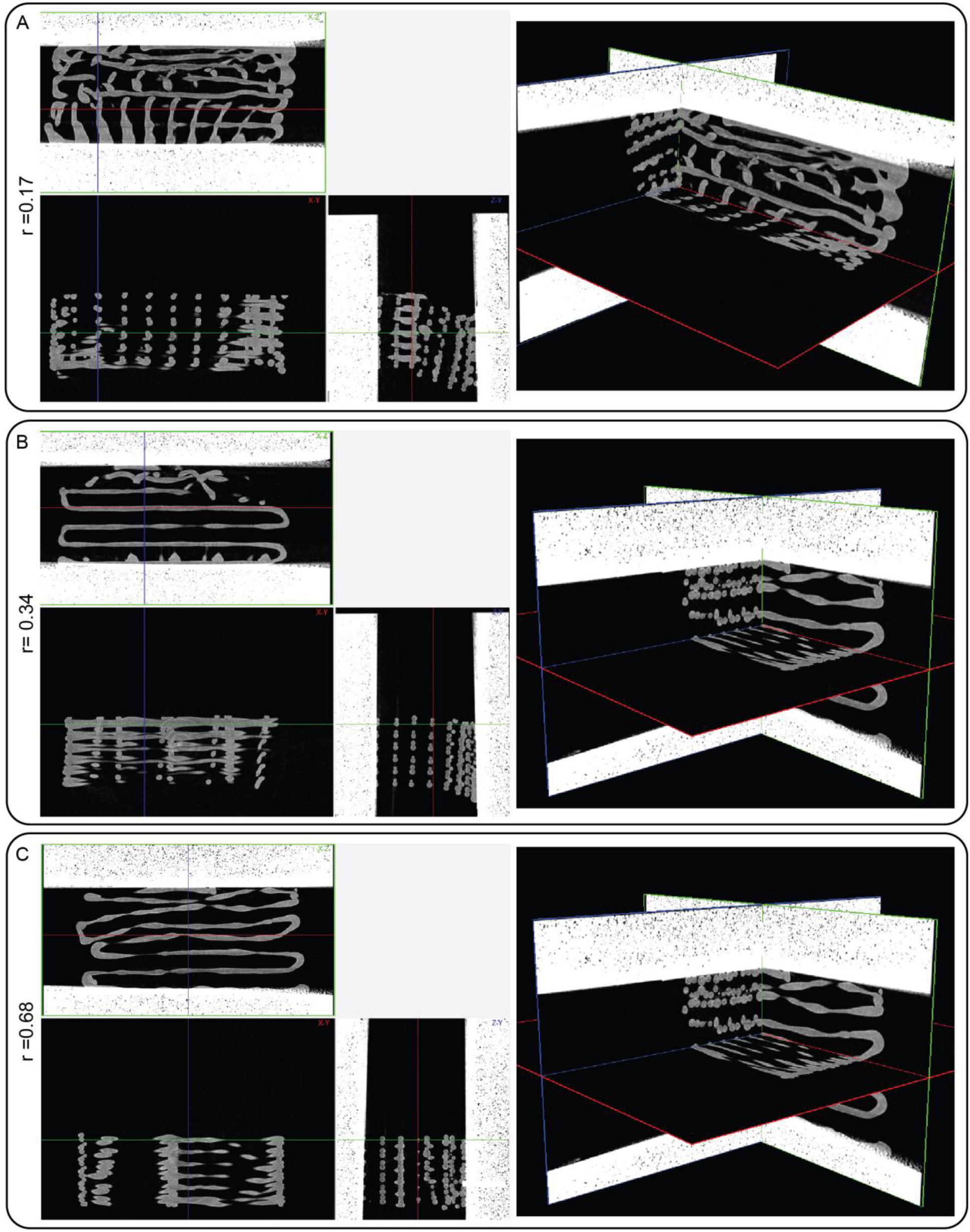
Micro-CT images after a compression test at 25% strain. Left panels reveal the cross-sections from different planes and the right panels reveal the 3D assembly of each plane. (A) r=0.17 design, (B) r=0.34 design and (C) the r=0.68 design.

**Figure S5.**
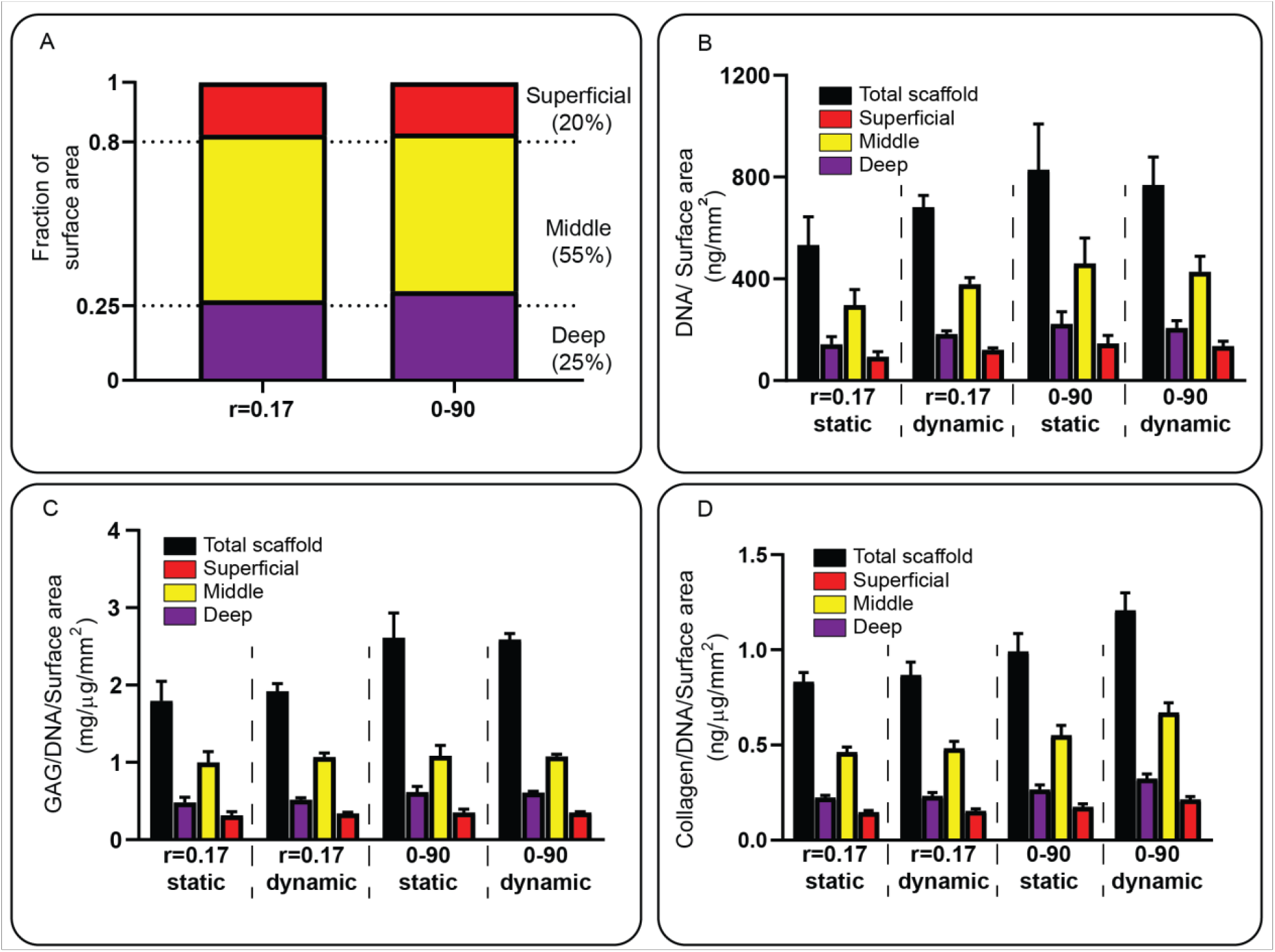
DNA,GAG and collagen analyses of the tested scaffolds after 28 days of culture. (A) Micro-CT analyses of the surface area in fraction of the surface area. (B) DNA content normalized against the surface area. (C) GAG per DNA content normalized against the surface area. (D) Collagen per DNA content normalized against the pore surface area. Each condition contained n=3 samples and values represent average ± standard deviation. * p<0.005.

**Figure S6.**
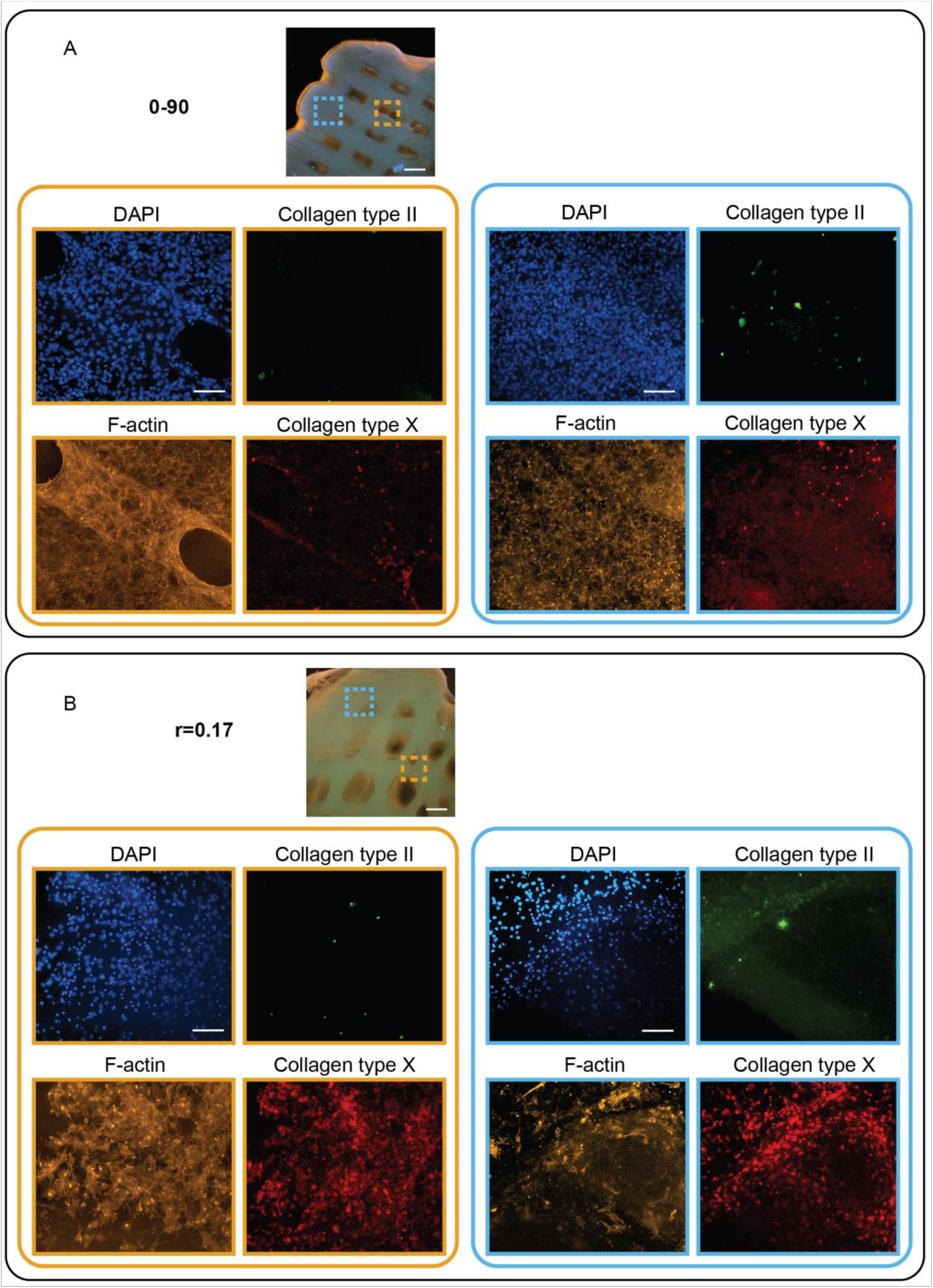
Cells after 28 days in static culture. On top of the panel an overview image of the scaffold. The bottom panel sets represent different areas in the scaffold. (A) 0-90 woodpile design. (B) r=0.17 hypotrochoidal design. Scale bars represent 500 μm in the overview panel and 200 μm in the close up.

**Figure S7.**
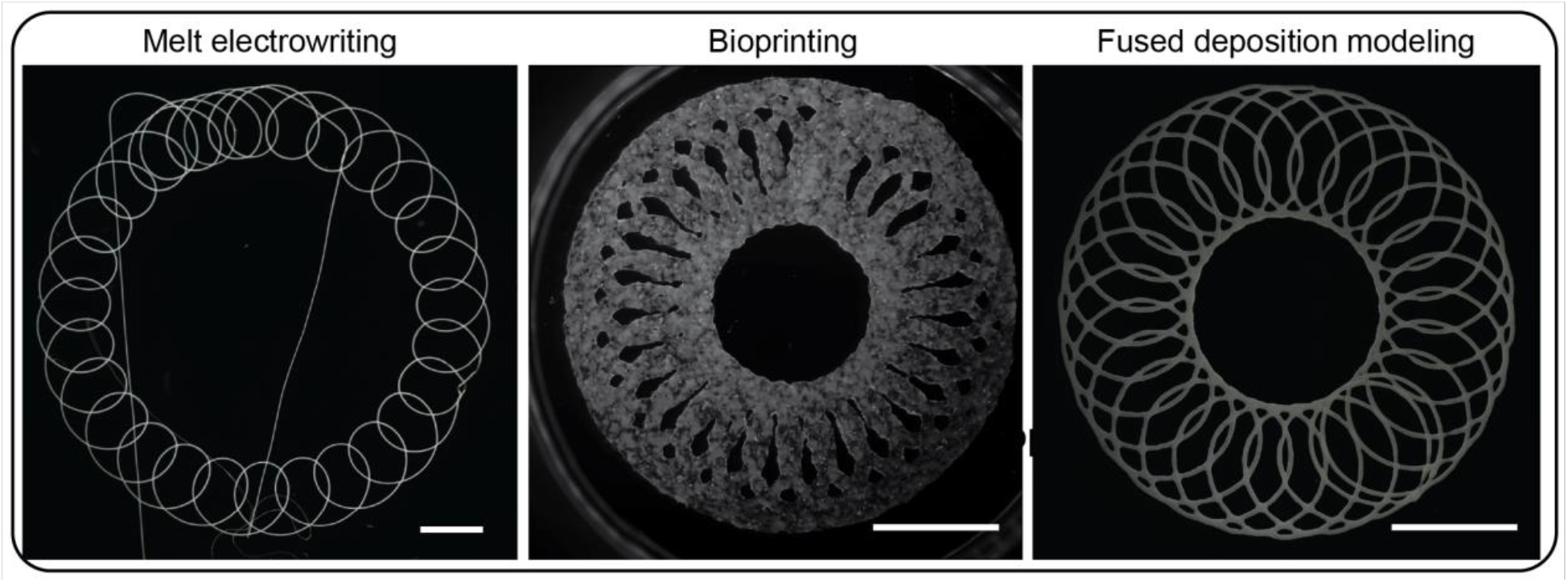
Hypotrochoidal designs fabricated with various techniques and printers. From left to right, Melt electrowntting, Bioprinting and Fused deposition modeling. With PCL Mn 45,000 as material for the melt electrowritten and fused deposition modeling scaffolds and Xanthan gum (7% w/v) as material for the bioprinted scaffolds. Scale bar represents 1000 μm.

## References

[1] J. Buckwalter, H. Mankin, Articular cartilage: part I, Journal of Bone and joint surgery 79(4) (1997) 600.

[2] A.J. Sophia Fox, A. Bedi, S.A. Rodeo, The basic science of articular cartilage: structure, composition, and function, Sports health 1(6) (2009) 461–468.

[3] L. Sharma, Osteoarthritis of the knee, New England Journal of Medicine 384(1) (2021) 51–59.

[4] D.J. Huey, J.C. Hu, K.A. Athanasiou, Unlike bone, cartilage regeneration remains elusive, Science 338(6109) (2012) 917–921.

[5] H. Muir, P. Bullough, A. Maroudas, The distribution of collagen in human articular cartilage with some of its physiological implications, The Journal of bone and joint surgery. British volume 52(3) (1970) 554–563.

[6] M. Ulrich-Vinther, M.D. Maloney, E.M. Schwarz, R. Rosier, R.J. O’Keefe, Articular cartilage biology, JAAOS-Journal of the American Academy of Orthopaedic Surgeons 11(6) (2003) 421–430.

[7] F. Guilak, D.L. Butler, S.A. Goldstein, Functional tissue engineering: the role of biomechanics in articular cartilage repair, Clinical Orthopaedics and Related Research^®^ 391 (2001) S295–S305.

[8] R.M. Aspden, D.W. Hukins, Collagen organization in articular cartilage, determined by X-ray diffraction, and its relationship to tissue function, Proceedings of the Royal Society of London. Series B, Biological sciences 212(1188) (1981) 299–304.

[9] R. Minns, F. Steven, The collagen fibril organization in human articular cartilage, Journal of anatomy 123(Pt 2) (1977) 437.

[10] T. Klein, B. Schumacher, T. Schmidt, K. Li, M. Voegtline, K. Masuda, E.-M. Thonar, R. Sah, Tissue engineering of stratified articular cartilage from chondrocyte subpopulations, Osteoarthritis and cartilage 11(8) (2003) 595–602.

[11] J.D. Lawrence, A catalog of special plane curves, Courier Corporation 2013.

[12] M. Bouthillier, Representations of Epitrochoids and Hypotrochoids, (2018).

[13] K. Nakanishi, H. Kojima, T. Watanabe, Trajectories of in-plane periodic solutions of tethered satellite system projected on van der Pol planes, Acta Astronautica 68(7-8) (2011) 1024–1030.

[14] C. Goldstein, J. Gray, J. Ritter, L’Europe mathématique/Mathematical Europe: Histoires, mythes, identités, Les Editions de la MSH1996.

[15] B. Glisic, New in Old: Simplified Equations for Linear-elastic Symmetric Arches and Insights on Their Behavior, Journal of the International Association for Shell and Spatial Structures 61(3) (2020) 227–240.

[16] C. Zhao, Y. Liu, P. Wang, M. Jiang, J. Zhou, X. Kong, Y. Chen, F. Jin, Wrapping and anchoring effects on CFRP strengthened reinforced concrete arches subjected to blast loads, Structural Concrete 22(4) (2021) 1913–1926.

[17] H.-J. Yen, C.-S. Tseng, S.-h. Hsu, C.-L. Tsai, Evaluation of chondrocyte growth in the highly porous scaffolds made by fused deposition manufacturing (FDM) filled with type II collagen, Biomedical microdevices 11(3) (2009) 615–624.

[18] M.E. Hoque, Y.L. Chuan, I. Pashby, Extrusion based rapid prototyping technique: an advanced platform for tissue engineering scaffold fabrication, Biopolymers 97(2) (2012) 83–93.

[19] L. Moroni, J.R. de Wijn, C.A. van Blitterswijk, 3D fiber-deposited scaffolds for tissue engineering: Influence of pores geometry and architecture on dynamic mechanical properties, Biomaterials 27(7) (2006) 974–985.

[20] K.-C. Hung, C.-S. Tseng, L.-G. Dai, S.-h. Hsu, Water-based polyurethane 3D printed scaffolds with controlled release function for customized cartilage tissue engineering, Biomaterials 83 (2016) 156–168.

[21] M.A. Shamekhi, H. Mirzadeh, H. Mahdavi, A. Rabiee, D. Mohebbi-Kalhori, M.B. Eslaminejad, Graphene oxide containing chitosan scaffolds for cartilage tissue engineering, International journal of biological macromolecules 127 (2019) 396–405.

[22] W. Kosorn, M. Sakulsumbat, P. Uppanan, P. Kaewkong, S. Chantaweroad, J. Jitsaard, K. Sitthiseripratip, W. Janvikul, PCL/PHBV blended three dimensional scaffolds fabricated by fused deposition modeling and responses of chondrocytes to the scaffolds, Journal of Biomedical Materials Research Part B: Applied Biomaterials 105(5) (2017) 1141–1150.

[23] S.M. Bittner, B.T. Smith, L. Diaz-Gomez, C.D. Hudgins, A.J. Melchiorri, D.W. Scott, J.P. Fisher, A.G. Mikos, Fabrication and mechanical characterization of 3D printed vertical uniform and gradient scaffolds for bone and osteochondral tissue engineering, Acta biomaterialia 90 (2019) 37–48.

[24] A. Di Luca, I. Lorenzo-Moldero, C. Mota, A. Lepedda, D. Auhl, C. Van Blitterswijk, L. Moroni, Tuning Cell Differentiation into a 3D Scaffold Presenting a Pore Shape Gradient for Osteochondral Regeneration, Advanced healthcare materials 5(14) (2016) 1753–63.

[25] J. Baena, G. Jiménez, E. López-Ruiz, C. Antich, C. Griñán-Lisón, M. Perán, P. Gálvez-Martín, J. Marchal, Volume-by-volume bioprinting of chondrocytes-alginate bioinks in high temperature thermoplastic scaffolds for cartilage regeneration, Experimental Biology and Medicine 244(1) (2019) 13–21.

[26] M. Uhl, C. Ihling, K. Allmann, J. Laubenberger, U. Tauer, C. Adler, M. Langer, Human articular cartilage: in vitro correlation of MRI and histologic findings, European radiology 8(7) (1998) 1123–1129.

[27] Q.L. Loh, C. Choong, Three-dimensional scaffolds for tissue engineering applications: role of porosity and pore size, Tissue Engineering Part B: Reviews 19(6) (2013) 485–502.

[28] R. Sayles, T. Thomas, J. Anderson, I. Haslock, A. Unsworth, Measurement of the surface microgeometry of articular cartilage, Journal of Biomechanics 12(4) (1979) 257–267.

[29] S.M. McNary, K.A. Athanasiou, A.H. Reddi, Transforming growth factor β-induced superficial zone protein accumulation in the surface zone of articular cartilage is dependent on the cytoskeleton, Tissue Engineering Part A 20(5-6) (2014) 921–929.

[30] S. Mohanty, L.B. Larsen, J. Trifol, P. Szabo, H.V.R. Burri, C. Canali, M. Dufva, J. Emnéus, A. Wolff, Fabrication of scalable and structured tissue engineering scaffolds using water dissolvable sacrificial 3D printed moulds, Materials science and engineering: C 55 (2015) 569–578.

[31] S. Roberts, J.P. Urban, H. Evans, S.M. Eisenstein, Transport properties of the human cartilage endplate in relation to its composition and calcification, Spine 21(4) (1996) 415–420.

[32] C.J. Little, N.K. Bawolin, X. Chen, Mechanical properties of natural cartilage and tissue-engineered constructs, Tissue Engineering Part B: Reviews 17(4) (2011) 213–227.

[33] D.E. Shepherd, B.B. Seedhom, The ‘instantaneous’ compressive modulus of human articular cartilage in joints of the lower limb, Rheumatology (Oxford, England) 38(2) (1999) 124–32.

[34] H. Chen, Y. Liu, C. Wang, A. Zhang, B. Chen, Q. Han, J. Wang, Design and properties of biomimetic irregular scaffolds for bone tissue engineering, Computers in Biology and Medicine 130 (2021) 104241.

[35] J.E. Jeffrey, D.W. Gregory, R.M. Aspden, Matrix damage and chondrocyte viability following a single impact load on articular-cartilage, Archives of biochemistry and biophysics 322(1) (1995) 87–96.

[36] L.T. Brody, Knee osteoarthritis: clinical connections to articular cartilage structure and function, Physical Therapy in Sport 16(4) (2015) 301–316.

[37] F. Tahmasebinia, Y. Ma, K. Joshua, S.M.E. Sepasgozar, Y. Yu, J. Li, S. Sepasgozar, F.A. Marroquin, Sustainable architecture creating arches using a bamboo grid shell structure: Numerical analysis and design, Sustainability 13(5) (2021) 2598.

[38] F. Malekipour, C. Whitton, D. Oetomo, P.V.S. Lee, Shock absorbing ability of articular cartilage and subchondral bone under impact compression, Journal of the mechanical behavior of biomedical materials 26 (2013) 127–135.

[39] B.M. Lawless, H. Sadeghi, D.K. Temple, H. Dhaliwal, D.M. Espino, D.W. Hukins, Viscoelasticity of articular cartilage: Analysing the effect of induced stress and the restraint of bone in a dynamic environment, Journal of the Mechanical Behavior of Biomedical Materials 75 (2017) 293–301.

[40] A. Weizel, T. Distler, D. Schneidereit, O. Friedrich, L. Bräuer, F. Paulsen, R. Detsch, A. Boccaccini, S. Budday, H. Seitz, Complex mechanical behavior of human articular cartilage and hydrogels for cartilage repair, Acta Biomaterialia 118 (2020) 113–128.

[41] A. Sharma, L.D. Wood, J.B. Richardson, S. Roberts, N.J. Kuiper, Glycosaminoglycan profiles of repair tissue formed following autologous chondrocyte implantation differ from control cartilage, Arthritis Research & Therapy 9(4) (2007) R79.

[42] T.E. Hardingham, A.J. Fosang, Proteoglycans: many forms and many functions, The FASEB journal 6(3) (1992) 861–870.

[43] P. Bansal, N.S. Joshi, V. Entezari, M. Grinstaff, B.D. Snyder, Contrast enhanced computed tomography can predict the glycosaminoglycan content and biomechanical properties of articular cartilage, Osteoarthritis and cartilage 18(2) (2010) 184–191.

[44] S. Treppo, H. Koepp, E.C. Quan, A.A. Cole, K.E. Kuettner, A.J. Grodzinsky, Comparison of biomechanical and biochemical properties of cartilage from human knee and ankle pairs, Journal of Orthopaedic Research 18(5) (2000) 739–748.

[45] G. Shen, The role of type X collagen in facilitating and regulating endochondral ossification of articular cartilage, Orthodontics & craniofacial research 8(1) (2005) 11–17.

[46] H. Akiyama, C. Shukunami, T. Nakamura, Y. Hiraki, Differential expressions of BMP family genes during chondrogenic differentiation of mouse ATDC5 cells, Cell structure and function 25(3) (2000) 195–204.

[47] C. Shukunami, K. Ishizeki, T. Atsumi, Y. Ohta, F. Suzuki, Y. Hiraki, Cellular hypertrophy and calcification of embryonal carcinoma-derived chondrogenic cell line ATDC5 in vitro, Journal of Bone and Mineral Research 12(8) (1997) 1174–1188.

[48] S.D. Waldman, C.G. Spiteri, M.D. Grynpas, R.M. Pilliar, R.A. Kandel, Long-term intermittent compressive stimulation improves the composition and mechanical properties of tissue-engineered cartilage, Tissue engineering 10(9-10) (2004) 1323–1331.

[49] C. Gaut, K. Sugaya, Critical review on the physical and mechanical factors involved in tissue engineering of cartilage, Regenerative medicine 10(5) (2015) 665–679.

[50] V.V. Meretoja, R.L. Dahlin, S. Wright, F.K. Kasper, A.G. Mikos, The effect of hypoxia on the chondrogenic differentiation of co-cultured articular chondrocytes and mesenchymal stem cells in scaffolds, Biomaterials 34(17) (2013) 4266–4273.

[51] D. MacKenna, S. Vaplon, A.D. McCULLOCH, Microstructural model of perimysial collagen fibers for resting myocardial mechanics during ventricular filling, American Journal of Physiology-Heart and Circulatory Physiology 273(3) (1997) H1576–H1586.

[52] M. Kääb, I. Ap Gwynn, H. Nötzli, Collagen fibre arrangement in the tibial plateau articular cartilage of man and other mammalian species, The Journal of Anatomy 193(1) (1998) 23–34.

[53] M.G. Patino, M.E. Neiders, S. Andreana, B. Noble, R.E. Cohen, Collagen: an overview, Implant dentistry 11(3) (2002) 280–285.

[54] J.P. Armstrong, E. Pchelintseva, S. Treumuth, C. Campanella, C. Meinert, T.J. Klein, D.W. Hutmacher, B.W. Drinkwater, M.M. Stevens, Tissue Engineering Cartilage with Deep Zone Cytoarchitecture by High-Resolution Acoustic Cell Patterning, Advanced healthcare materials (2022) 2200481.

[55] G. Hochleitner, E. Fürsattel, R. Giesa, J. Groll, H.W. Schmidt, P.D. Dalton, Melt electrowriting of thermoplastic elastomers, Macromolecular rapid communications 39(10) (2018) 1800055.

[56] V.I. Sikavitsas, J.S. Temenoff, A.G. Mikos, Biomaterials and bone mechanotransduction, Biomaterials 22(19) (2001) 2581–2593.

[57] Q. Fu, E. Saiz, A.P. Tomsia, Direct ink writing of highly porous and strong glass scaffolds for load-bearing bone defects repair and regeneration, Acta Biomaterialia 7(10) (2011) 3547–3554.

